# Single cell proteomics characterization of bone marrow hematopoiesis with distinct Ras pathway lesions

**DOI:** 10.1101/2023.12.20.572584

**Authors:** Laila Karra, Anna-Marie Finger, Lauren Shechtman, Milana Krush, Rita Meng-Yao Huang, Morgan Prinz, Iliana Tennvooren, Kriti Bahl, Lisiena Hysienaj, Paulina G. Gonzalez, Alexis J. Combes, Hugo Gonzalez, Rafael J Argüello, Matthew H. Spitzer, Jeroen P. Roose

## Abstract

Normal hematopoiesis requires constant prolific production of different blood cell lineages by multipotent hematopoietic stem cells (HSC). Stem- and progenitor- cells need to balance dormancy with proliferation. How genetic alterations impact frequency, lineage potential, and metabolism of HSC is largely unknown. Here, we compared induced expression of KRAS^G12D^ or RasGRP1 to normal hematopoiesis. At low-resolution, both Ras pathway lesions result in skewing towards myeloid lineages. Single-cell resolution CyTOF proteomics unmasked an expansion of HSC- and progenitor- compartments for RasGRP1, contrasted by a depletion for KRAS^G12D^. SCENITH™ quantitates protein synthesis with single-cell precision and corroborated that immature cells display low metabolic SCENITH™ rates. Both RasGRP1 and KRAS^G12D^ elevated mean SCENITH™ signals in immature cells. However, RasGRP1-overexpressing stem cells retain a metabolically quiescent cell-fraction, whereas this fraction diminishes for KRAS^G12D^. Our temporal single cell proteomics and metabolomics datasets provide a resource of mechanistic insights into altered hematopoiesis at single cell resolution.

## INTRODUCTION

The skin, intestine, and hematopoietic system are examples of tissues with dynamic turnover of mature cells dying off or leaving the tissue, therefore necessitating prolific production of daughter cells from ever self-renewing stem cells (Fuchs, 2009). Multi-dimensional, proteomic profiling of hematopoiesis by single-cell mass cytometry (CyTOF) has delineated a developmental continuum, born out of multipotent hematopoietic stem cells (HSC) (Bendall et al., 2011). The combined need for life-long self-renewal and high production of terminally differentiated cells is secured via a homeostatic balance with deeply dormant HSC that harbor the highest stem cell activity and cycling HSC that produce offspring (**Figure 1A**) (Rossi et al., 2012; Wilson et al., 2009). Disruption of this homeostatic balance of dormant and cycling HSC state can lead to bone marrow failure and is also observed in some hematopoietic malignancies (Orford and Scadden, 2008).

**Figure 1:**
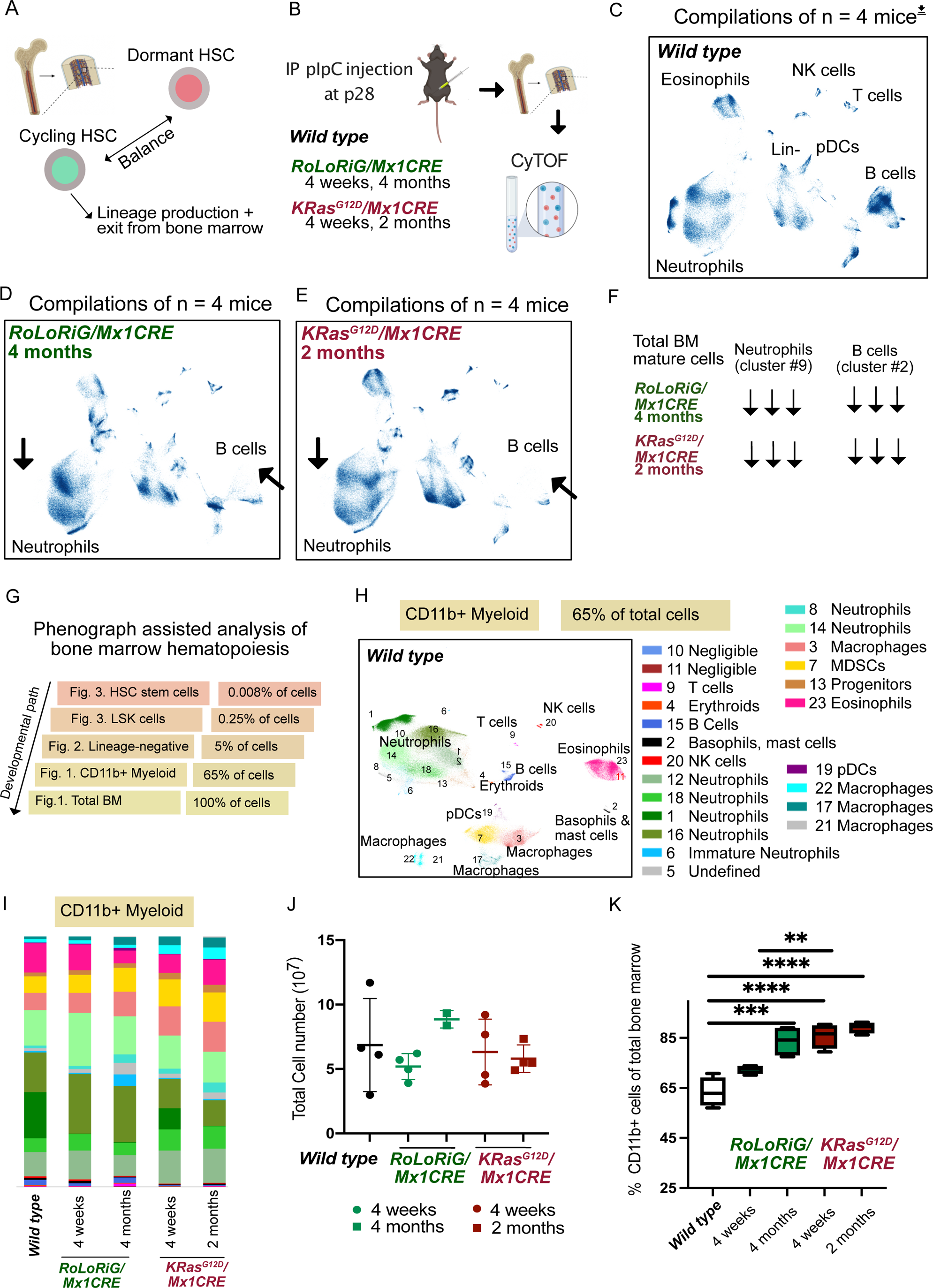
High dimensional analysis of induced, aberrant Ras signals in the mature bone marrow compartment. (A) A balance of dormant and cycling HSC. (B) pIpC injection of the indicated genetic mouse models induces the desired Ras pathway lesions. (C, D, E) UMAP representation of cell composition in wild type BM, RoLoRiG/Mx1CRE total BM, and KRASG12D/Mx1CRE total BM. UMAPs are compilations of n = 4 mice at the indicated post-pIpC injection time. (F) Summary of main points for total BM analysis. (G) Strategy of using single cell resolution CyTOF to investigate stem and progenitor cells. Frequencies of cell types are calculated from wild type mice, based on our CyTOF data. (H) PhenoGraph analysis revealing 23 CD11b-positive clusters in BM of wild type mice, compilation of n = 4 mice. Cluster #10 and 11 are identified by PhenoGraph but datapoints are negligible and #5 is undefined (Supplemental Table S2). (I) Relative abundance of the 23 clusters, visualizing overall changes in the CD11b-positive compartment. (J) Total bone marrow cell counts in the 20 individual mice. (K) CD11b-positive cells as a percentage of total bone marrow cells for the indicated experimental groups.**p<0.01 ***p<0.001 ****p<0.0001.

Acute Lymphoblastic Leukemia (ALL) is one of the leading causes of cancer death in persons under 20 years old. Challenging characteristics of ALL that impact leukemia patients include the uncertain developmental origin (Good et al., 2018; Zhang et al., 2012), plasticity of ALL cells (Alexander et al., 2018), and heterogeneity in the hematopoietic ALL composition (Bendall et al., 2014; Bendall et al., 2011; Kleppe et al., 2017). Specific genetic backgrounds or gene mutations in mouse models can promote enhanced cycling activity of HSC with an initial increase but a subsequent loss of HSC numbers (Orford and Scadden, 2008). This implies that leukemic cells that carry mutated oncogenes must somehow retain the self-renewing stem cell properties required for a durable growth advantage. However, gaining mechanistic insights into this balance has been hampered due the scarcity of HSCs.

Somatic mutations in *RAS* genes, a family of small GTPases, or lesions in other genes that aberrantly elevate Ras signals are highly prevalent in blood cancers, particularly juvenile myelomoncytic leukemia and other myeloid malignancies (Bowen et al., 2005; Liu et al., 2017; Schubbert et al., 2007; von Lintig et al., 2000; Ward et al., 2012). While less common in acute lymphoblastic leukemia ALL), *RAS* gene mutations are enriched in early T precursor ALL (Zhang et al., 2012). In addition, over-expression of the Ras activator *RasGRP1* (Ras guanine nucleotide releasing protein 1) frequently occurs in T-ALL (Hartzell et al., 2013) and is mutually exclusive with somatic mutations in *RAS* (Dail et al., 2010). The biochemical impacts of this collection of genetic lesions, such as the amplitude and duration of Ras signals or the cycling patterns through the RasGDP (off) – RasGTP (on) states, are well-documented (**Supplemental Figure S1A**) (Ksionda et al., 2013; Kulhanek et al., 2021; Mues and Roose, 2016). Here, we focused on two distinct Ras pathway lesions, induced expression of KRas^G12D^ and the over-expression of RasGRP1. These lesions were chosen as representative examples of different Ras pathway disruptions, aimed at investigating the alterations in the hematopoietic continuum and stem- and progenitor- cell characteristics within the bone marrow. We introduced these single genetic alterations in bone marrow (BM) cells by capitalizing on the *Mx1CRE* allele and injection of polyinosinic-polycytidylic acid (pIpC) (Kuhn et al., 1995) (**Figure 1B**).

To create a comprehensive resource detailing alterations in hematopoiesis, we conducted a CyTOF single-cell proteomics analysis to characterize the dynamic changes occurring in the hematopoietic continuum (See **Supplemental Figure S1B**). We find that, while both KRas^G12D^ and RasGRP1 expression led to a skewing towards the myeloid lineage, with CD11b-positive cells dominating the bone marrow over time, their effects on the stem cell compartment were markedly distinct. Our proteomics studies revealed that KRas^G12D^ resulted in a depletion of the HSC- and LSK- (Lineage negative, Sca1- and cKit-positive; containing HSC) stem- and progenitor- cell populations, whereas RasGRP1 promoted an expansion of these rare cell subsets. Furthermore, these two aberrant Ras signals had distinct impacts on the lineage commitment of HSCs to multipotent progenitors, diverging not only from *wild type* cells but also from each other.

Mechanistically, we utilized single-cell resolution of translational activity by ribosomes using SCENITH™ (for Single-Cell ENergetIc metabolism by profiling Translation inHibition) (Arguello et al., 2020) in conjunction with spectral flow (Hysenaj et al., 2023). The resolution provided by spectral flow revealed that Stem- and progenitor- cells exhibit lower metabolic SCENITH™ rates compared to mature, lineage-positive hematopoietic cells, corroborating the notion that Stem- and progenitor- cells are metabolically more quiescent. The lineage-negative compartments with Stem- and progenitor- cells demonstrated elevated SCENITH™ values for both the KRas^G12D^ and RasGRP1 Ras pathway lesions. However, the fraction of cells that displays a low protein translation rate in the *wild type* LSK compartment is diminished in KRas^G12D^ LSK yet retained in LSK that overexpress RasGRP1. HSCs with either Ras pathway lesion display increased SCENITH™ values, but for KRas^G12D^ this is paired with a further reduction in HSC fraction of the already depleted LSK compartment.

Our single cell proteomic studies provide a comprehensive resource for understanding how single, induced genetic alterations impact the frequency, lineage potential, and ribosome translation rates of rare stem cells within the bone marrow.

## STAR**★**METHODS

### CONTACT FOR REAGENT AND RESOURCE SHARING

Further information and requests for resources and reagents should be directed to and will be fulfilled by the Lead Contact, Jeroen Roose.

### EXPERIMENTAL MODEL

#### Mice

All *wild type* mice were derived from the C57BL6 strain purchased from Jackson lab as control animals. Engineered *“KRas^G12D^*mice” inducibly express a mutated glycine to aspartic acid amino acid at position 12 (G12D) via a LoxP-STOP-LoxP cassette from the endogenous *KRas* locus (Jackson et al., 2001). *KRas^G12D^* mice were purchased from Jackson and cross-bred with *Mx1CRE* mice, expressing the IFN-α/β-inducible Mx-1 promoter (Kuhn et al., 1995) kindly provided by Dr. Emmanuel Passague. We previously generated *RoLoRiG* (Rosa26, LoxP-STOP-LoxP, RasGRP1, ires, GFP) mice that, when crossed to *Mx1CRE* mice and injected with pIpC, allow for RasGRP1 overexpression in conjunction with GFP (Karra et al., 2020). Such mice live a normal lifespan but develop a mild splenomegaly (Karra et al., 2020). Animals were culled at 4 weeks, 2 months, and 4 months post intra-peritoneal polyinosinic-polycytidylic acid (pIpC; 1 μg/250 μl/mouse, Sigma-Aldrich cat. no. P0913-50MG) at approximately p56 days of age. All mice were housed and treated in accordance with the guidelines at the Institutional Animal Care and Use Committee guidelines of the UCSF, protocol AN195708-01C.

### METHOD DETAILS

#### Single cell suspension generation from bone marrow

*RoLoRiG/Mx1CRE* and *KRAS^G12D^/Mx1CRE* mice were injected with pIpC and euthanized after

4 weeks and 4 months for *RoLoRiG/Mx1CRE* mice, and 4 weeks and 2 months for *KRAS^G12D^/Mx1CRE* mice. Tibia and femurs of mice were collected and cleaned from flesh. Bones were then crushed using mortar and pestle in Hank’s Balanced Salt Solution (HBSS; UCSF cell culture facility) containing 2% of FBS. Cells were filtered through a 70 μm strainer. Red blood cell lysis was performed for all tissues with ACK buffer (150 mM NH_4_Cl, 10 mM KHCO_3_, 0.1 mM Na_2_EDTA at pH 7.2-7.4). Bone marrow (BM) cells were washed once with HBSS, then loaded onto a Histopaque-1119 (Sigma-Aldrich, San Luis, MO, US) density gradient and centrifuged at 1500 rpm with no break for 5 min to separate bone debris from cells. All cells were then counted and kept on ice for CyTOF or SCENITH™ experiments.

#### CyTOF-Preparation of cells for staining

After obtaining the primary cells from the bone marrow they were prepped as previously described (Spitzer et al., 2017). Briefly, bone marrow cells were washed in PBS + 5mM EDTA at 4°C and centrifuged at 500g for 5 min. Cells were resuspended at 1:1 with PBS with 5mM EDTA and 100μM Cisplatin (Sigma-Aldrich, San Luis, MO, US) for 60s before quenching 1:1 with PBS with 0.5% BSA and 5mM EDTA to determine viability as previously described (Spitzer et al., 2015). Cells were centrifuged at 500g for 5 min at 4°C and resuspended in PBS with 0.5% BSA and 5mM EDTA at a density between 1–10*10^6^ cells/ml. Cells were fixed for 10 min at RT using 1.6% PFA (Fisher Scientific, Hampton, New Hampshire) and then kept at −80°C.

#### Mass Cytometry Antibodies

All mass cytometry antibodies and concentrations used for analysis can be found in **Supplemental Table S1**. Primary conjugates of mass cytometry antibodies were prepared using the MaxPAR antibody conjugation kit (Fluidigm) according to the manufacturer’s recommended protocol. After labeling, antibodies were diluted in Candor PBS Antibody Stabilization solution (Candor Bioscience) supplemented with 0.02% NaN3 to between 0.1 and 0.3 mg ml−1 and stored long term at 4°C. Each antibody clone and lot was titrated to optimal staining concentrations using primary murine samples.

#### Mass-Tag Cellular Barcoding

Mass-tag cellular barcoding was performed as previously described; Each sample is positive for three tags and negative for three other tags to create a unique “barcode” (Zunder et al., 2015). Briefly, 1*10^6^ cells from each animal were barcoded with distinct combinations of stable Pd isotopes chelated by isothiocyanobenzyl-EDTA in 0.02% saponin in PBS. Samples from any given tissue from one mouse per treatment group were barcoded together, with at least 3 biological replicates per treatment group across different plates. Cells were washed two times in PBS with 0.5% BSA and 0.02% NaN3 and pooled into a single FACS tube (BD Biosciences). After data collection, each condition was deconvoluted using a single-cell debarcoding algorithm (Zunder et al., 2015).

#### Mass Cytometry Staining and Measurement

Cells were resuspended in cell-staining media (PBS with 0.5% BSA and 0.02% NaN3) and metal-labeled antibodies against CD16/32 were added at 20μg/ml for 5 min at RT on a shaker to block Fc receptors. Surface marker antibodies (**Supplemental Table S1**) were then added, yielding 500uL final reaction volumes and stained at room temperature for 30min at RT on a shaker. Following staining, cells were washed twice with cell-staining media and then permeabilized with 4°C methanol for at 10 min at 4°C. Cells were then washed twice in cell-staining media to remove remaining methanol, and then stained with intracellular antibodies in 500 μL for 30 min at RT on a shaker. Cells were washed twice in cell-staining media then stained with 1 mL of 1:4000 191/193Ir DNA intercalator (Fluidigm) diluted in PBS with 1.6% PFA overnight. Cells were then washed once with cell-staining media and then two times with double-deionized (dd)H20. Care was taken to assure buffers preceding analysis were not contaminated with metals in the mass range above 100 Da. Mass cytometry samples were diluted in ddH2O containing bead standards (see below) to approximately 10^6^ cells per mL and then analyzed on a CyTOFTM 2 mass cytometer (Fluidigm) equilibrated with ddH2O. We analyzed 1–5*10^5^ cells per animal and per time point, consistent with generally accepted practices in the field.

#### Mass Cytometry Bead Standard Data Normalization

Data normalization was performed as previously described (Spitzer et al., 2017). Briefly, just before analysis, the stained and intercalated cell pellet was resuspended in freshly prepared Cell Acquisition Solution containing the bead standard at a concentration ranging between 1 and 2*10^4^ beads/ml. The mixture of beads and cells were filtered through a filter cap FACS tubes (BD Biosciences) before analysis. All mass cytometry files were normalized together using the mass cytometry data normalization algorithm (Finck et al., 2013), which uses the intensity values of a sliding window of these bead standards to correct for instrument fluctuations over time and between samples.

#### SCENITH™ Metabolic Assay

SCENITH™ (Arguello et al., 2020). relies on (1) incorporation of puromycin in polypeptides at the ribosome, (2) the fact polypeptide synthesis is energetically costly, and (3) a novel anti-puro monoclonal antibody that allows for assessment of the puromycin incorporation level with individual cell resolution. SCENITH™ kit containing all reagents (including anti-Puromycin clone R4743L-E8) clone and protocols were obtained (www.scenith.com) and the metabolic profile was analyzed using the method as previously described (Arguello et al., 2020). Briefly, live cells were plated in quadruplets at a concentration of (1.30*10^5^ cells in 180.5ul media/well) in a U-bottom 96 well plate immediately post extraction. After equilibrating to room temperature, 9.5μL of Control (DMSO), 2-Deoxy-D-Glucose (DG; [100mM] Sigma-Aldrich Cat. No. D6134), Oligomycin (Oligo; [1 μM Sigma-Aldrich Cat. No.75351), or equal parts DG and Oligo at the final concentrations listed were added for 30-minute incubation at 37°C. Puromycin at was then added in media and plates were incubated for an additional 1 hour at 37°C. Following puro treatment, cells were washed with cold PBS to stop the reaction and centrifuged at 1500rpm, 5 minutes at RT. Cells were then stained with 100ul of 1:5000 dilution of viability dye (Zombie NIR, ThermoFisher) in PBS. and incubated in the dark for 20 minutes at RT. After washing with 100uL HBSS buffer/2% FBS, cells were stained with a mix of antibodies for the different stem cell subsets. Cells were washed once and centrifuged at 1500 rpm for 5 minutes. Cells were then fixed/permed with 100uL fixation solution (FoxP3 Transcription Factor Staining Buffer Kit; Tonbo Biosciences cat. no. TNB-0607-KIT) for 20 minutes in the dark. Cells were then washed and resuspended in 100ul of HBSS/2% FBS overnight. On the second day of preparation, cells were centrifuged at 2500rpm for 2 minutes and washed twice with permeabilization buffer using a 1:10 dilution with ddH20. Cells were blocked with 100ul per well 2% NRS (Normal Rat Serum; Thermo Fisher cat. No.10710C) in permeabilization buffer for 20 minutes at RT. Without washing, 20uL of anti-puromycin monoclonal antibody (Arguello et al., 2020)(produced in the lab of Dr. Rafael Arguello; Clone R4743L-E8 conjugated with Alexa Fluor 647 or with Alexa Fluor 488) was added and incubated for 30 minutes in the dark. At this step additional wells were stained with anti-KI67 or matching isotype control antibody [5ug/ml]. Cells were washed once with permeabilization buffer and once with FACS buffer by centrifuging at 2500rpm for 2 minutes and discarding supernatant. To prepare for flow cytometry, cells were merged from all replicates into one well containing 150-200ul and transferred into microFACS tubes. Samples were acquired on Cytek Aurora at 400000 events per sample. All data were analyzed with FlowJo software (Tree Star).

#### Spectral Flow Data Visualization by Dimensionality Reduction

Unmixed and compensated spectral flow data was exported from SpectroFlo software (Cytek Biosciences) and imported to FlowJo. Manual gating was performed on cellular events > singlets > live cells (Zombie NIR^−^) > lineage negative cells (CD3^−^, CD4^−^, CD5^−^, CD11b^−^, CD8^−^, CD45R^−^, Ly76/TER-119^−^), for which FCS files were exported. Data transformation, quality control, and dimensionality reduction was a performed essentially as described (den Braanker et al., 2021). Briefly, flow cytometry data was imported to R using the flowCore package (Hahne et al., 2009) and was normalized using the arcsinh cofactor transformation method of the flowVS package (Azad et al., 2016). Automated quality control of transformed data was performed with the PeacoQC package (Emmaneel et al., 2022) before flow data was converted to a SingleCellExperiment object in R. Subsequently, tSNE plots of spectral flow data were generated by dimensionality reduction and visualization using Seurat and scater toolkits (Hao et al., 2021; McCarthy et al., 2017). During clustering, forward and side scatter, lineage markers, Zombie NIR, and autofluorescence were excluded for tSNE generation. FCS files for spectral flow experiments and computational pipelines used in this study are available upon request.

#### Unsupervised clustering and compositional distance

Each cell subset—all cells, CD11b+ cells, lin-cells, and LSK cells—was clustered separately using the PhenoGraph clustering algorithm (Levine et al., 2015) as implemented in the ‘cytofkit’ package in R. Standard settings were utilized (k=15, minimum distance=0.1). Cluster frequencies were calculated as a percent of total cells within that subset. Aitchison distances were calculated between the composition of the cluster frequencies of each individual mouse compared to the cluster frequencies of a random wild-type mouse and were calculated in R.

#### Data Visualization

Dimensionality reduction was performed using the UMAP algorithm implemented in Python. In the UMAP plot, each individual cell was colored according to its cluster or was colored according to its expression of denoted protein of interest. Heat maps of the resulting cluster frequencies and the hierarchical clustering of the heat maps were generated with the Seaborn package in Python. The log2 fold change heat maps were calculated as the log2 of the fold change in cluster frequencies between the indicated two groups.

### QUANTIFICATION AND STATISTICAL ANALYSIS

Statistical significance of results was calculated using Prism 9 (GraphPad Software). Analysis was performed using one-way ANOVA with tukey-post test or a student’s *t* test when comparing two groups. A p value <0.05 was considered significant and is indicated with an asterisk.

### MATERIALS AVAILABILITY

This study did not generate new unique reagents.

## RESULTS

### Single cell proteomics characterization of total bone marrow of *wild type, RoLoRiG/Mx1CRE,* and *KRAS^G12D^/Mx1CRE* mice

In normal cells, Ras cycles between RasGDP (*off*) and RasGTP (*on*) states at regulated rates. Ras’ intrinsic GTPase activity normally secures efficient conversion to the *off* state, enhanced by RasGAPs (Ras GTPase Activating Proteins) (**Supplemental Figure S1A**) (Ksionda et al., 2013; Kulhanek et al., 2021). RasGRP, SOS, and other RasGEFs (Ras guanine nucleotide exchange factors) catalyze nucleotide exchange on Ras, which then rebinds free GTP or GDP in the cell. Nucleotide exchange increases Ras-GTP levels due to the much higher intracellular concentrations of GTP than GDP. Overexpression of RasGRP1 results in abnormally increased cycling between RasGTP/RasGDP, whereas KRas^G12D^ is trapped in the GTP-bound *on* state due to a reduced rate of intrinsic GTP hydrolysis and resistance to GTPase activating proteins (GAPs) (**Supplemental Figure S1A**) (Mues and Roose, 2016). Here, we characterized the impact of these two distinct aberrant Ras signals on hematopoiesis in a temporal manner by comparing the bone marrow (BM) compartment of control *wild type* mice to *RoLoRiG/Mx1CRE* and *KRAS^G12D^/Mx1CRE* mice with single cell resolution, using CyTOF (**Figure 1B**).

Single BM cell suspensions were counted, fixed, stained, and analyzed in parallel using a mass cytometry panel for 40 surface markers, leaving 6 channels for mass-tag cellular barcoding (Zunder et al., 2015) for the twenty mice in the five experimental groups (**Supplemental Figure S1C** and **Supplemental Table S1**). The Uniform Manifold Approximation and Projection (UMAP) algorithm was used to project cellular BM compositions (**Supplemental Figure S1D**), generating a standard of total *wild type* BM (**Figure 1C**). Using such high-level surveys, major changes in mature BM populations were readily detected, like a gradual loss of B cells in both the *RoLoRiG/Mx1CRE* and *KRAS^G12D^/Mx1CRE* models (**Figures 1D**, **1E**, and **Supplemental Figures S1E** and **S1F**).

We next performed PhenoGraph analysis (Levine et al., 2015) to cluster cell populations into groups based on protein expression values extracted from all 40 surface markers, each of which was color-coded in the UMAP visualization. Phenograph analysis on total BM identified 26 cell clusters (**Supplemental Figures S2A-C** and **S3**) and for each of the 26 clusters we extracted quantitative marker expression data to generate an unbiased measure of cell identity, depicted in heat maps (**Supplemental Figure S4**). We performed hierarchical clustering of the heat maps with the Seaborn package in Python, which provides a resource on common and rare populations in the total bone marrow (**Supplemental Figure S5**), and alterations in the 26 clusters for total bone marrow in which the two distinct Ras pathway lesions were induced (**Supplemental Figure S6**). In **Supplemental Figures S6B** and throughout the manuscript, we utilized the *wild type* genotype as a reference to compare dynamically altered BM composition from *RoLoRiG/Mx1CRE* and *KRAS^G12D^/Mx1CRE* mice.

Using these high-level surveys, the total BM of *RoLoRiG/Mx1CRE* and *KRAS^G12D^/Mx1CRE* mice diverges from *wild type* over time, evidenced by the calculated compositional distance (**Supplemental Figure S2D**) with notable disappearance of B cell and neutrophil clusters (**Figure 1F**). At this level of resolution *RoLoRiG/Mx1CRE* and *KRAS^G12D^/Mx1CRE* total BM compartments display only modest unique features between the two (**Supplemental Figure S6C**).

We next performed single cell proteomics characterizations of hematopoiesis with PhenoGraph analyses, working in a reverse-developmental path from mature, lineage-committed cells to stem cells, capitalizing on the single cell resolution and a stepwise focus on the very rare LSK- and HSC-compartments (**Figure 1G**).

### Ras pathway lesions drive increased production of myeloid lineages with relatively intact developmental maturation

*KRas^G12D^/Mx1CRE* mice develop an aggressive myeloproliferative disease (MPD) (Braun et al., 2004; Chan et al., 2004), whereas the *RoLoRiG/Mx1CRE* model displays a mild myeloproliferation (Karra et al., 2020). We therefore zoomed-in on the CD11b-positive, myeloid compartment. PhenoGraph analysis defined 23 clusters of cell populations within the CD11b-positive dataset from *wild type*, *RoLoRiG/Mx1CRE,* and *KRasG12D/Mx1CRE* BM (**Figure 1H** and **Supplemental Figures S7A-C**). The relative abundance of the 23 CD11b-positive cell clusters (**Figure 1I**) and the color-coded Log2-fold changes in cell cluster frequencies (**Supplemental Figure S7D**) revealed that the overall patterns for these 23 clusters were very similar between the three genetic models. Thus, the developmental paths of cells maturing in the myeloid lineage (**Supplemental Figure S1B**) are mostly intact in *RoLoRiG/Mx1CRE* and *KRasG12D/Mx1CRE* BM. However, while the total number of BM cells was similar for all five experimental set-ups (**Figure 1J**), the fractions of CD11b-positive cells were strongly increased with both Ras pathway lesions; CD11b-positive cells dominate the bone marrow composition, already after 4 weeks in KRasG12D/Mx1CRE mice and after 4 months in RoLoRiG/Mx1CRE animals (**Figure 1K**).

### Unique CMP and contrasting LSK progenitor cell populations in *RoLoRiG/Mx1CRE* and *KRASG12D/Mx1CRE* mice

Upon *KRas^G12D^* induction, LSK cells (Lineage negative, Sca1- and cKit-positive; containing HSC) lose their quiescence through unknown mechanisms and decrease in numbers (Sabnis et al., 2009). By contrast, *RoLoRiG/Mx1CRE* stem or progenitor cells appear more fit in native hematopoiesis settings (Karra et al., 2020), but the LSK- and HSC-compartments in *RoLoRiG/Mx1CRE* mice have remained unexplored. To understand the origin of remodeled hematopoiesis resulting from distinct Ras pathway lesions, we employed PhenoGraph to analyze progenitor cells in *RoLoRiG/Mx1CRE* and *KRas^G12D^/Mx1CRE* mice. Computational gating on the lineage negative (Lin-) compartment that excludes the majority of BM cells with mature cell markers CD11b, CD3, CD4, CD5, CD8, TER119, and B220 provides the required resolution to investigate stem- and progenitor- cell populations (**Supplemental Figure S8A**).

PhenoGraph analysis resulted in 20 clusters of stem- and progenitor- cell populations in the lineage-negative compartment, including CMP (Common Myeloid Progenitors) and the LSK population that contains HSC (**Figure 2A** and **Supplemental Table S3**). The lineage negative-compartments of *RoLoRiG/Mx1CRE* and *KRAS^G12D^/Mx1CRE* mice significantly parted from *wild type* mice but also from each other (**Figures 2A-2D** and **Supplemental Figure S8B**), with unique emergence of undefined populations, such as cluster #6 in *KRAS^G12D^/Mx1CRE* mice and cluster #11 in *RoLoRiG/Mx1CRE* and *KRAS^G12D^/Mx1CRE* mice, compared to *wild type* counterparts (See full surface marker expression profiles in **Supplemental Table S3**). The calculated compositional distance enumerates the divergence in the lineage-negative compartment (**Supplemental Figure S8C**).

**Figure 2:**
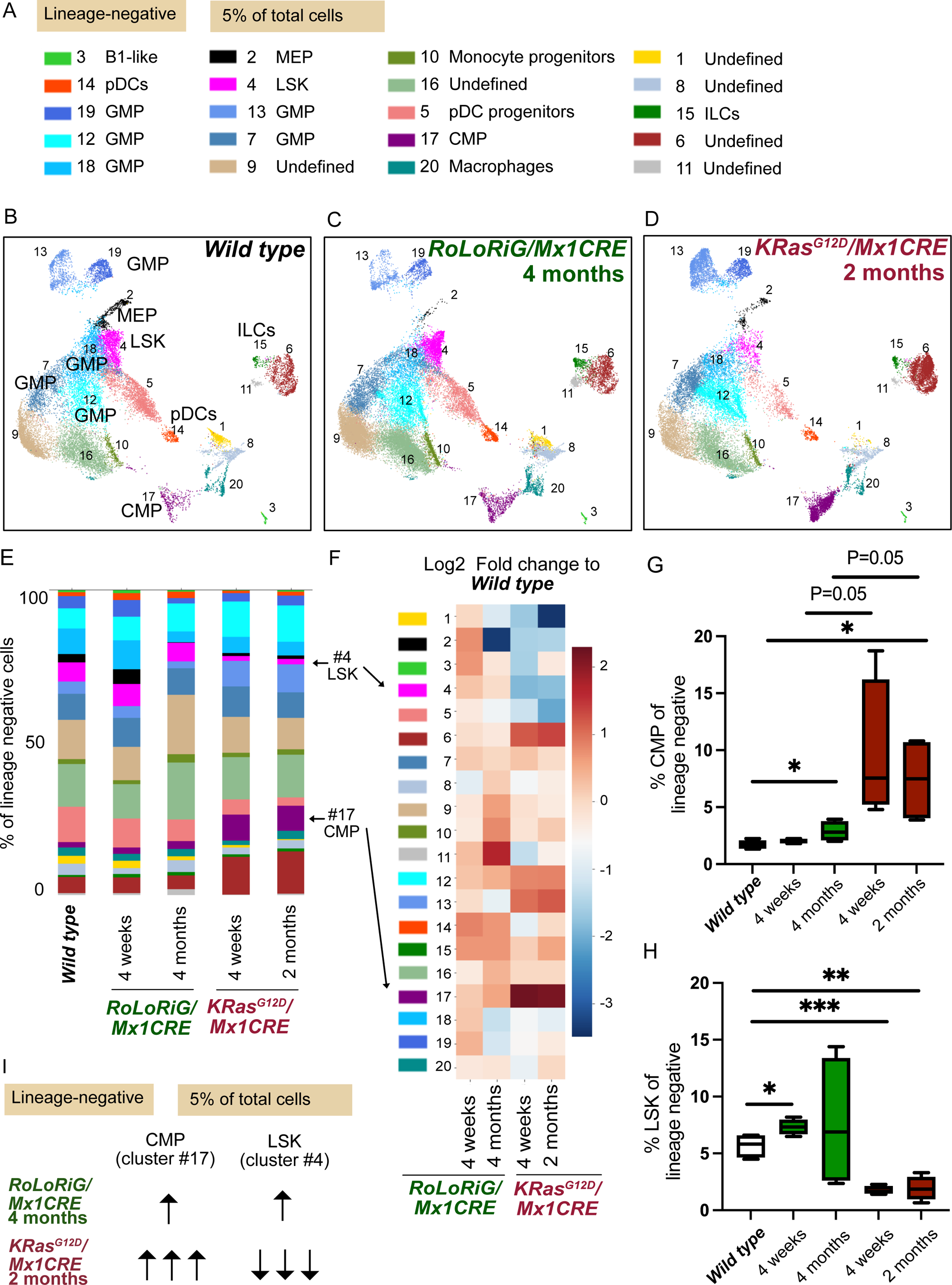
Distinct Ras pathway lesions drive opposing immature BM compartment features. (A) PhenoGraph analysis of lineage-negative cells reveal 20 cell clusters in BM (B-D) PhenoGraph analysis of lineage-negative compartment for *wild type* BM, *RoLoRiG/Mx1CRE* total BM, and *KRAS^G12D^/Mx1CRE* total BM. UMAPs are compilations of n = 4 mice at the indicated post-pIpC injection time. (E) Relative abundance of the 20 assigned, lineage marker-negative cell clusters in the bone marrow. Note the frequencies of LSK cells (#4) and CMP (#17). (F) Color-coded visualization of Log2 fold changes in cell cluster frequencies of the lineage-negative compartment in *RoLoRiG/Mx1CRE* and *KRAS^G12D^/Mx1CRE* BM, compared to *wild type*. (G, H) Frequencies of CMP and LSK stem cells as a percentage of lineage-negative bone marrow cells. *p<0.05 **p<0.01 ***p<0.001. (I) Summary of main points for analysis of lineage-negative compartment.

We next focused on two specific, lineage-negative populations that provide deeper understanding of the impacts of KRas^G12D^ or RasGRP1 overexpression on native hematopoiesis *in vivo*. First, CMP cluster #17 strongly and rapidly expanded upon pIpC-induced KRas^G12D^ expression, but much more modestly and delayed in *RoLoRiG/Mx1CRE* mice (**Figure 2A-2F** and **Supplemental Figure S8B**). CMP fuel the CD11b-positive myeloid cell lineages in normal BM maturation (**Supplemental Figure S1B**). The respective increases in CMP in the *RoLoRiG/Mx1CRE* and *KRas^G12D^/Mx1CRE* models (**Figure 2G**) closely mirrored the magnitude of elevated CD11b-positive cell fractions (**Figure 1K**), implying that the aberrantly expanded CMP generally follow maturation to CD11b-positive myeloid cell lineages in the bone marrow of both models.

Remarkably, the LSK cluster #4, which contains HSC, revealed opposing phenotypes in response to the two aberrant Ras signals. In agreement with originally reports using traditional, lower resolution flow cytometry (Sabnis et al., 2009), LSK cell percentages decreased in *KRas^G12D^/Mx1CRE* mice, compared to *wild type* (**Figures 2D-F**, **2H**, and **Supplemental Figure S8B**). By contrast, the LSK compartment from *RoLoRiG/Mx1CRE* mice was expanded (**Figures 2C** and **2H**).

### Dynamic remodeling of the *RoLoRiG/Mx1CRE* LSK compartment with increased HSC population

Using the field’s stem cell annotation in which MPPs 1-4 arise from HSCs (Wilson et al., 2008) (**Figure 3A** and **Supplemental Figure S8D**), we characterized 10 clusters in the *wild type* LSK compartment with high resolution, encompassing a #8 CLP-like cluster, #4 MPP3-like, #5 MPP4-like, and two unidentified cell clusters #7 and #10 (**Figure 3B** and **Supplemental Table S4**). The reduced cellularity of the *KRAS^G12D^/Mx1CRE* LSK compartment (**Figure 2I**) was even more evident at this higher resolution (**Figure 3C**, **Supplemental Figure S8E**) and paired with a relative shunting of the *KRAS^G12D^/Mx1CRE* LSK subpopulations towards #9 MPP2 and away from #2 MPP4 (**Figures 3E-G**, **3I** and **Supplemental Figure S9**). Note that we calculated the relative abundance of MPP2 and HSC as a percentage of LSK cells in Figure 3 and that LSK cells are reduced in *KRAS^G12D^/Mx1CRE* mice (**Figure 2H**).

**Figure 3:**
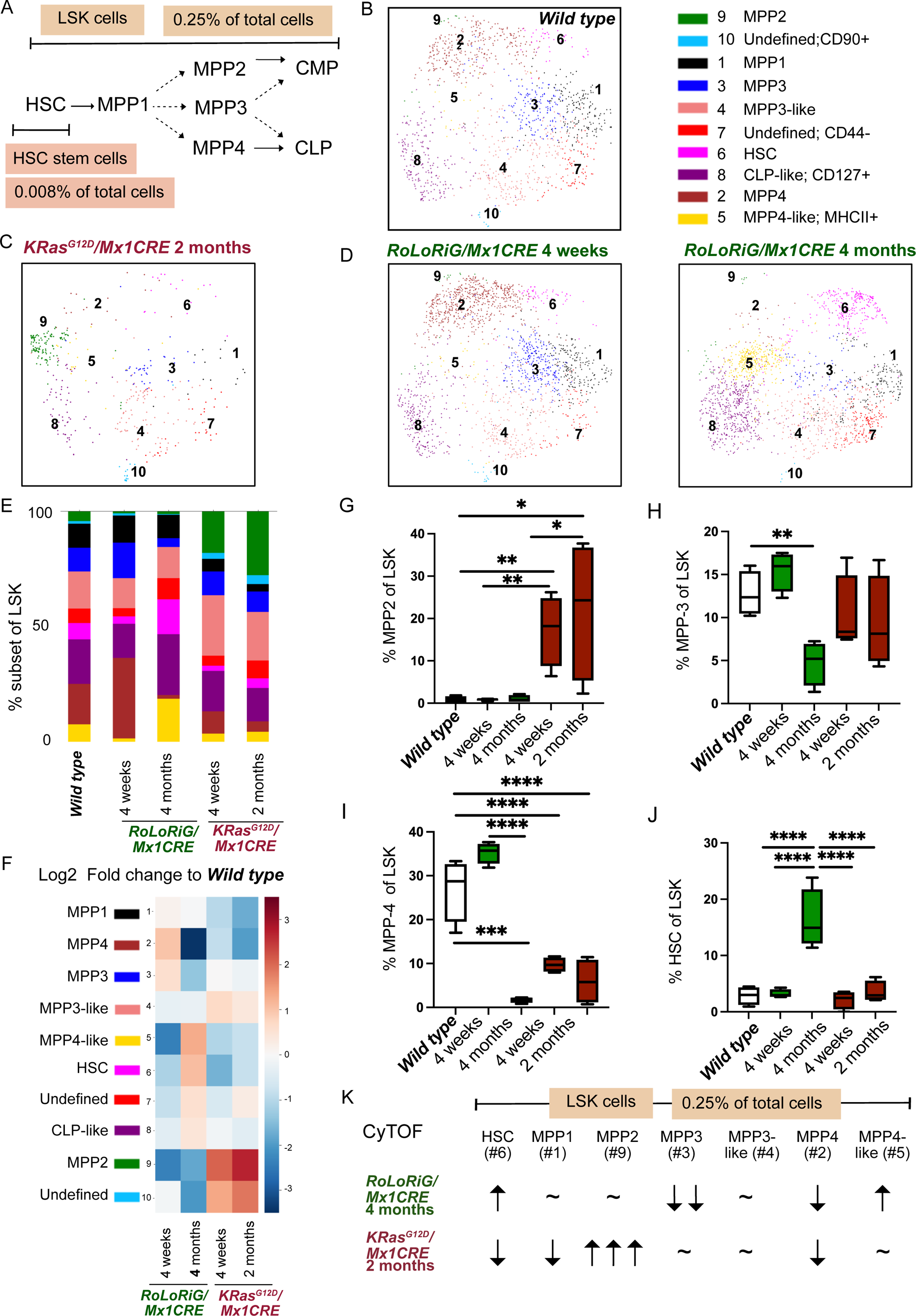
High-resolution analysis of hematopoietic stem cell compartments of *wild type, RoLoRiG/Mx1CRE* and *KRAS^G12D^/Mx1CRE* mice. (A) Schematic of HSC and MPP 1-4 annotation (Wilson et al., 2008), applied in this study. (B) PhenoGraph analysis of the *wild type* LSK compartment reveals 10 clusters: HSC, MPP1-4, CLP-like, MPP3-like, MPP4-like and two undefined populations. Compilation of n = 4 mice (C) As in 3B, but for *KRAS^G12D^/Mx1CRE*, 2 months post pIpC injection. (D) As in 3B, but for *RoLoRiG/Mx1CRE* mice. (E) Relative abundance of the 10 clusters in the LSK compartment. (F) Color-coded visualization of Log2 fold changes in LSK cell cluster frequencies of *RoLoRiG/Mx1CRE* and *KRAS^G12D^/Mx1CRE* BM, compared to *wild type*. (G-J) Frequencies of MPP2, MPP3, MPP4, and HSC cells as a percentage of LSK bone marrow cells for the indicated experimental groups. *p<0.05 **p<0.01 ***p<0.001 ****p<0.0001. (K) Summary of main points for analysis of LSK compartment by CyTOF.

By contrast, the increased cellularity in the *RoLoRiG/Mx1CRE* LSK compartment (**Figure 2H**) occurred through a dynamic remodeling of this compartment that occurs between 4 weeks and 4 months post-pIpC injection (**Figures 3D-3H**). Initially, increases in #2 MPP4 and #3 MPP3 appeared, but these trends switched to reduced fractions of these populations at 4 months (**Figures 3F**, **3H**, and **3I**). At 4 months, the *RoLoRiG/Mx1CRE* LSK compartment was characterized by strong increases in #5 MPP4-like (**Supplemental Figure S9B**) and a striking accumulation of #6 HSC (**Figure 3J** and **Supplemental Figure S9F**).

Altogether, these findings show that HSC in mice with the distinct Ras pathway lesions follow uniquely rewired developmental paths through the MPP and MPP-like subsets (**Figure 3K** and **Supplemental Figure S9C-F**). Shared between the models is the aberrantly increased CMP population (**Figure 2G**) that results in CD11b+ myeloid cells taking over the bone marrow compartment (**Figure 1K**). Strikingly, the HSC pool in *KRAS^G12D^/Mx1CRE* mice is depleted while it is enhanced in *RoLoRiG/Mx1CRE* mice (**Figure 3K** and **Supplemental Figure S9C-F**).

### Measuring hematopoietic cell metabolism at single cell resolution

Hematopoietic Stem- and progenitor- cells appear to require finely tuned rates of protein synthesis (Signer et al., 2014), a cell biological process that is linked to available ATP and cell metabolism. Potent oncogenes, like *KRAS^G12D^*, result in metabolic rewiring cells, such as the Warburg effect (Mukhopadhyay et al., 2021). We reported that RasGRP1 signals to a metabolic mTOR pathway in T cells (Myers et al., 2019) and that *RoLoRiG/Mx1CRE* BM cells display increased fitness *ex vivo* in CFU assays in media-poor conditions (Karra et al., 2020). We therefore postulated that distinct metabolic features may underlie the opposing phenotype of HSC in the *RoLoRiG/Mx1CRE* and *KRAS^G12D^/Mx1CRE* models. The scarcity of LSC and HSC in the bone marrow (**Supplemental Figure S8F**) necessitates an experimental platform with single cell resolution and we next analyzed hematopoietic progenitor cell metabolism, directly *ex vivo* using SCENITH™ (Arguello et al., 2020).

In short, SCENITH™ relies on measuring changes of protein synthesis levels in response to metabolic inhibitors and allows the assessment of multiple cell types in parallel at single cell resolution (**Figure 4A**). As such, SCENITH™ measures the global levels of translation, which represents a proxy for cellular metabolic activity. As illustration of SCENITH™ analysis, four monoclonal T cell leukemia lines demonstrated unique but uniform puromycin staining patterns and this SCENITH™ signal collapsed when the *in vitro* puromycin exposure was performed in conjunction with metabolic pathway inhibitors (**Figure 4B**). By contrast, *wild type* hematopoietic cells reveal heterogeneous SCENITH™ signals (**Figure 4C**). As expected, addition of 2-Deoxy-D-glucose (2DG - to inhibit glycolysis) and oligomycin (OG - to inhibit mitochondrial ATP synthase) collapsed the SCENITH™ signal (**Figure 4C**).

**Figure 4:**
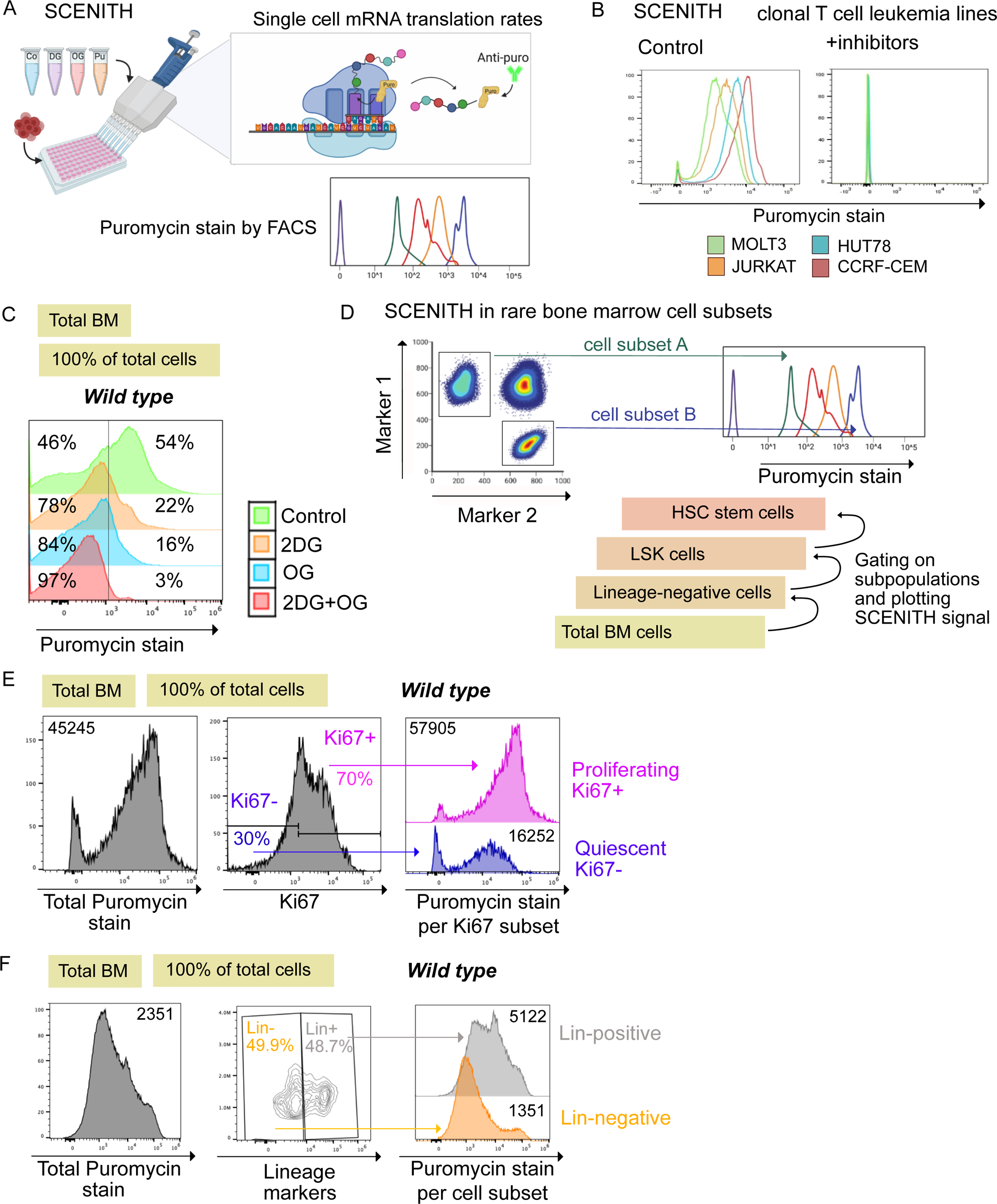
Low SCENITH™ ribosome activity in lineage-negative bone marrow cells. (A) SCENITH™ analysis of protein synthesis levels at the ribosome. Incorporated Puromycin is quantitatively measured. (B) Distinct uniform SCENITH™ signals in four monoclonal T cell leukemia cell lines. (C) Representative example of SCENITH™ pattern for wild type total bone marrow cells and effect of metabolic inhibitors on SCENITH™ signals. 2DG = 2-Deoxy-D-glucose, glycolysis inhibitor. OG = Oligomycin, inhibiting ATP Synthase. Percentages indicate cell fractions on either side of the arbitrarily set SCENITH™ signal cut-off. (D) Concept of combining SCENITH™ analysis with Spectral Flow to obtain cell subset-specific protein synthesis levels. (E) Comparisons of SCENITH™ in Ki67-negative and -positive fractions in total wild type bone marrow. Numbers in the histograms indicate the mean fluorescent intensity (MFI) for the anti-PURO antibody signal. (F) SCENITH™ in *wild type* lineage-negative Stem- and progenitor- cells, compared to lineage-positive, mature cells. Figures 4B, C, E, and F are representative example of at least 3 independent biological experiments.

Capitalizing on SCENITH™ as a flow cytometry-based method, we combined SCENITH™ with cell subset-defining marker panels to measure hematopoietic cell metabolism at single cell resolution (**Figure 4D**); Proliferating, Ki67-positive or lineage-positive BM cells demonstrated substantially higher protein synthesis levels than Ki67-negative and lineage-negative BM cells (**Figures 4E and 4F**). Thus, quiescent hematopoietic cells and Stem- and progenitor- cells have modest levels of metabolic activity, as assessed by SCENITH™, in agreement with reported low levels of protein synthesis (Signer et al., 2014).

### Metabolic assessment of hematopoietic cells with aberrant Ras signals

SCENITH™ allowed us to obtain mechanistic insights into the *KRAS^G12D^/Mx1CRE* and *RoLoRiG/Mx1CRE* models, analyzed four to eight weeks post pIpC injection. Note that we always compared our models with Ras pathway lesions to *wild type* controls performed in parallel on the same day.

Mean SCENITH™ values were increased for *KRAS^G12D^/Mx1CRE* and *RoLoRiG/Mx1CRE* total BM (**Figures 5A-5C**). The distribution of the SCENITH™ signals amongst individual cells depicted in histograms also allowed for enumeration of the fraction of *KRAS^G12D^/Mx1CRE* or *RoLoRiG/Mx1CRE* cell at low versus high levels of metabolic activity, displaying low or high puromycin incorporation signals (**Supplemental Figures S10A** and **S10B**). In *RoLoRiG/Mx1CRE* total BM the puromycin-low fraction was indistinguishable from *wild type* but the fraction of puromycin-high cells was increased (**Figure 5D**), whereas *KRAS^G12D^/Mx1CRE* total BM demonstrated a relative loss of the fraction of puromycin-low cells (**Figure 5E**).

**Figure 5:**
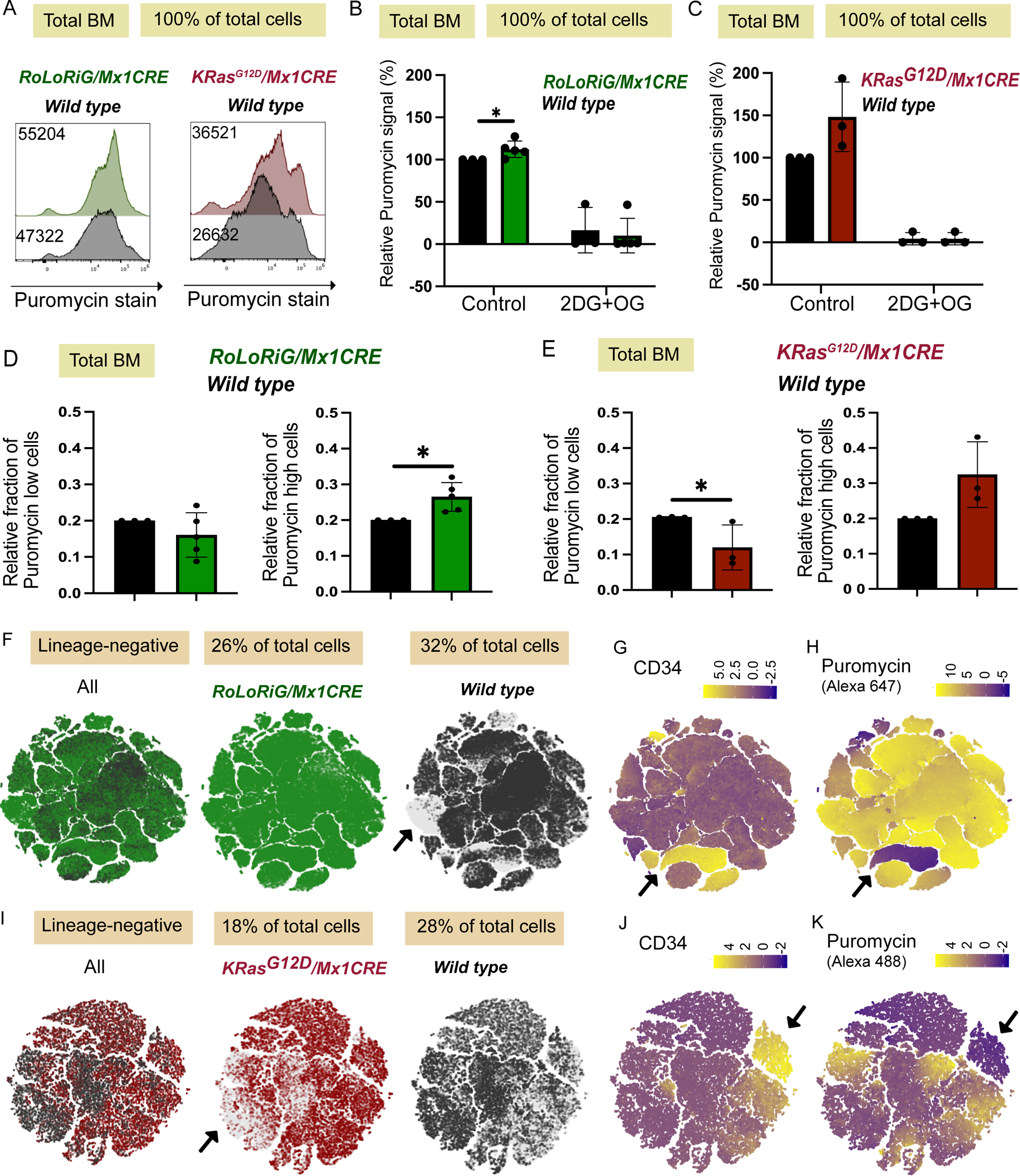
Metabolic comparisons of BM compartments with Ras pathway lesions. (A) Representative histograms of SCENITH™ analysis for *KRAS^G12D^/Mx1CRE* and *RoLoRiG/Mx1CRE* total bone marrow, compared to *wild type*. Numbers in the histograms indicate the mean fluorescent intensity (MFI) for the anti-PURO antibody signal. Representative example of at least 3 independent biological experiments. (B, C) Percentage relative SCENITH™ signal for *RoLoRiG/Mx1CRE* (n=5), compared to *wild type* (n=3). *p<0.05. In each individual experiment, comparing wild type to aberrant Ras signals, wild type control was arbitrarily set at 100%, allowing for comparisons across separate experiments. (C) for KRASG12D/Mx1CRE (n = 3) with three different wild type controls in which we used a Alexa488-coupled anti-PURO antibody. (D) Bar graphs of fractions of Puromycin-low and Puromycin-high total bone marrow cells for RoLoRiG/Mx1CRE and wild type, setting the wild type arbitrarily at 0.2 following the gating depicted in Supplemental Figure S10A. *p<0.05. (E) As in 5D, but for KRASG12D/Mx1CRE and wild type total BM. (F-H) t-SNE plots representing cell composition, CD34 expression levels, and SCENITH™ levels for lineage negative BM cells from individual RoLoRiG/Mx1CRE and wild type mice. For two additional pairs of mice, See Supplemental figure S11. (I-K) As in 5F-H, but for the KRASG12D/Mx1CRE and wild type lineage-negative compartment.

We subsequently combined SCENITH™ with Spectral Flow analysis so that cell subset-defining antibodies (**Resource Table**) enable metabolic assessment of rare, lineage-negative stem- and progenitor- populations (**Supplemental Figures S10C** and **S10D**). We first processed the SCENITH™/Spectral Flow data by defining gates for lineage-negative cells in *wild type* and copying these same gates onto the Ras pathway lesion models (per individual experiment, see **Supplemental Figures S10E** and **S11A**). t-SNE plots, projecting all lineage-negative cell data in 2D, pointed to differences in composition of the lineage-negative compartment (**Figures 5F and 5I**); differences that we had established with CyTOF analysis in Figure 2. Superimposing expression data for the CD34 stem cell marker and SCENITH™ signals, demarcated early stem and progenitor cells on the t-SNE lots and revealed that these cells have low metabolic activity (**Figures 5F-K** and **Supplemental Figures S11**).

### SCENITH™ analysis of lineage negative- and LSK-compartments with distinct Ras lesions

Next, we quantitated protein synthesis levels in all lineage-negative cells. Both *KRAS^G12D^/Mx1CRE* and *RoLoRiG/Mx1CRE* revealed increases in mean protein synthesis rates compared to *wild type* controls (**Figures 6A** and **6B**). The puromycin-low fraction was similar in *wild type* and *RoLoRiG/Mx1CRE* lineage-negative cells but decreased in the *KRAS^G12D^/Mx1CRE* lineage-negative compartment (**Figures 6C** and **6D**). Thus lineage-negative *KRAS^G12D^/Mx1CRE* cells increase ribosome translation at the expense of a metabolically quiescent fraction, whereas *RoLoRiG/Mx1CRE* lineage-negative counterparts can drive higher mean translational rates while still preserving a metabolically quiescent fraction.

**Figure 6:**
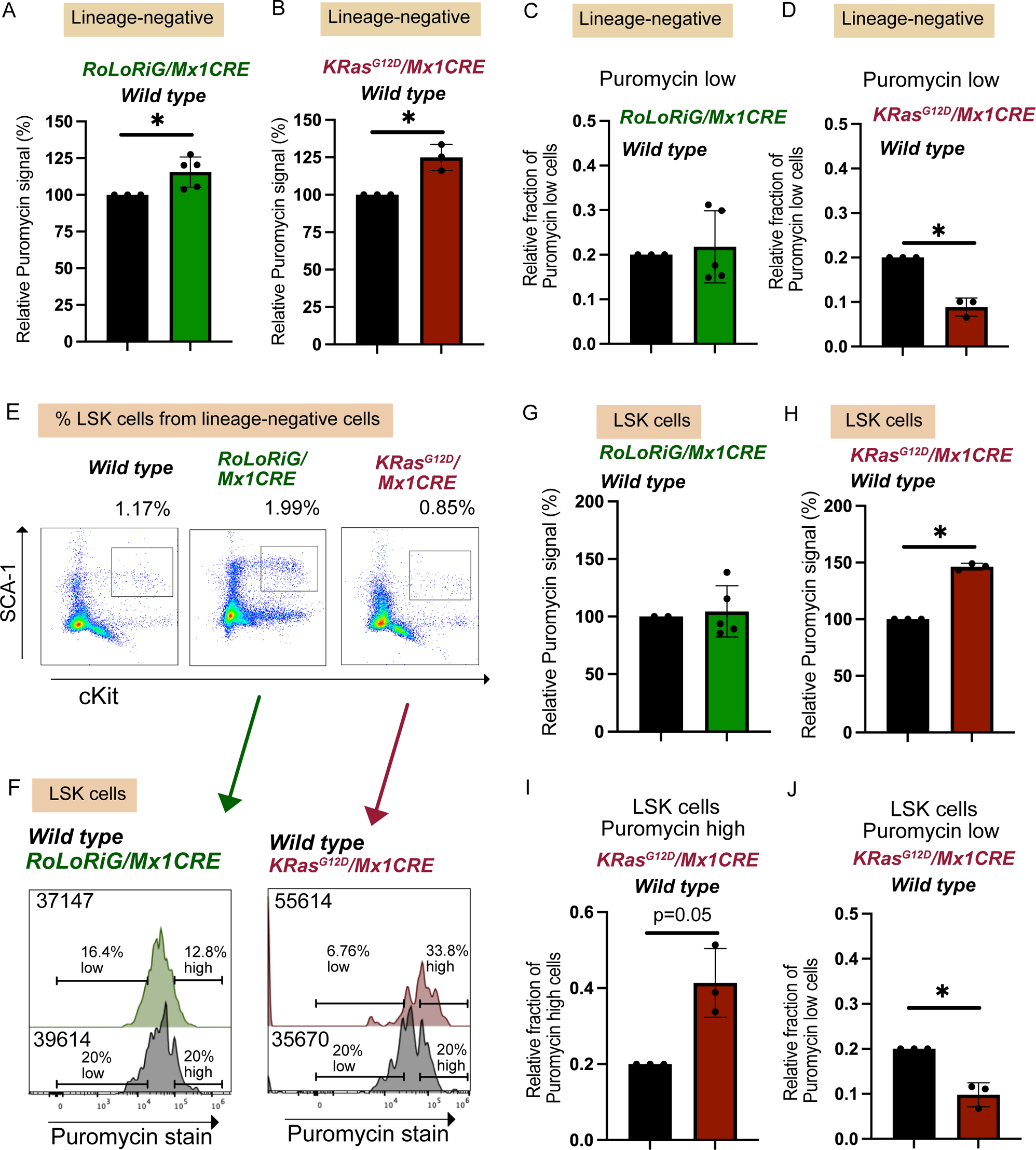
SCENITH™ analysis of the lineage negative – and LSK-bone marrow compartments. (A, B) Bar graphs representing relative SCENITH™ signals in lineage-negative cells from (A) *RoLoRiG/Mx1CRE* BM (n=5) and *wild type* (n = 3) or (B) *KRAS^G12D^/Mx1CRE* (n = 3) and *wild type* (n=3) BM. *Wild type* control was arbitrarily set at 100% for both A and B, *p<0.05. (C, D) Fractions of Puromycin-low lineage-negative cells for the indicated genotypes, setting *wild type* arbitrarily at 0.2 as in 5D. (E) Representative spectral flow dot plots of cKit- and SCA-1-positive LSK cells and their frequency. (F) Representative SCENITH™ histograms of LSK cells from the indicated mouse models. Large numbers depict the MFI for the anti-PURO antibody signal, small numbers the fraction of cells with low or high puromycin signal. 6E and 6F are representative examples of at least 3 independent biological experiments. (G, H) Bar graphs representing relative SCENITH™ signals in LSK cells from (G) *RoLoRiG/Mx1CRE* (n=5) and *wild type* (n=3) or (H) *KRAS^G12D^/Mx1CRE* (n = 3) and *wild type* (n = 3) BM. *Wild type* control was arbitrarily set at 100%, *p<0.05. (I, J) Bar graphs of fractions of Puromycin-high and Puromycin-low LSK cells for *KRAS^G12D^/Mx1CRE* and *wild type,* setting the *wild type* arbitrarily at 0.2. *p<0.05.

Deeper analysis of the lineage-negative population provided LSK-specific characterizations. Spectral flow confirmed our CyTOF data from Figure 3 with an enhanced *RoLoRiG/Mx1CRE* LSK compartment and depleted *KRAS^G12D^/Mx1CRE* LSK population (**Figure 6E** and **Supplemental Figure S10E**). *RoLoRiG/Mx1CRE* LSK cells displayed metabolic activities that resembled *wild type* counterparts, both in terms of mean SCENITH™ signals and fraction in the low- and high-puromycin cell subsets (**Figures 6F, 6G**, and data not shown). In sharp contrast, the reduced proportion of *KRAS^G12D^/Mx1CRE* cells that remained in the LSK gate showed increased mean translation levels (**Figure 6F** and **6H**). Further, this metabolic increase for *KRAS^G12D^/Mx1CRE* was paired with an expanded puromycin-high LSK cell fraction and loss of the puromycin-low LSK cell fraction (**Figure 6I and 6J**). Thus, the KRAS^G12D^ lesion is incompatible with maintaining LSK compartment that is relatively dormant in terms of protein synthesis levels.

### MPP- and HSC- stem cell compartments with distinct Ras pathway lesions

Lastly, we interrogated the rare, multipotent MPP and the HSC stem cell compartments in *KRAS^G12D^/Mx1CRE* and *RoLoRiG/Mx1CRE* models, analyzed four to eight weeks post pIpC injection. At this timepoint, we did not yet observe a remodeling of the MPP progenitors that overexpress RasGRP1 (**Figure 7A**), compared to the 4-month timepoint in Figure 3K. However, we did already note altered protein translation with relative increases in mean SCENITH™ signals for MPP1 and decreases for MPP2 and MPP4 (**Figure 7B**). *KRAS^G12D^/Mx1CRE* multipotent progenitors assessed by spectral flow showed skewing toward MPP2 and away from MPP3 and MPP4 (**Figure 7C**), like our CyTOF analyses (**Figure 3G-I**). The *KRAS^G12D^/Mx1CRE* progenitors that remained in the MPP1 through 4 subsets as defined by the indicated markers (**Supplemental Figure S12**) displayed a wide spread of SCENITH™ signals (**Figure 7D**).

**Figure 7:**
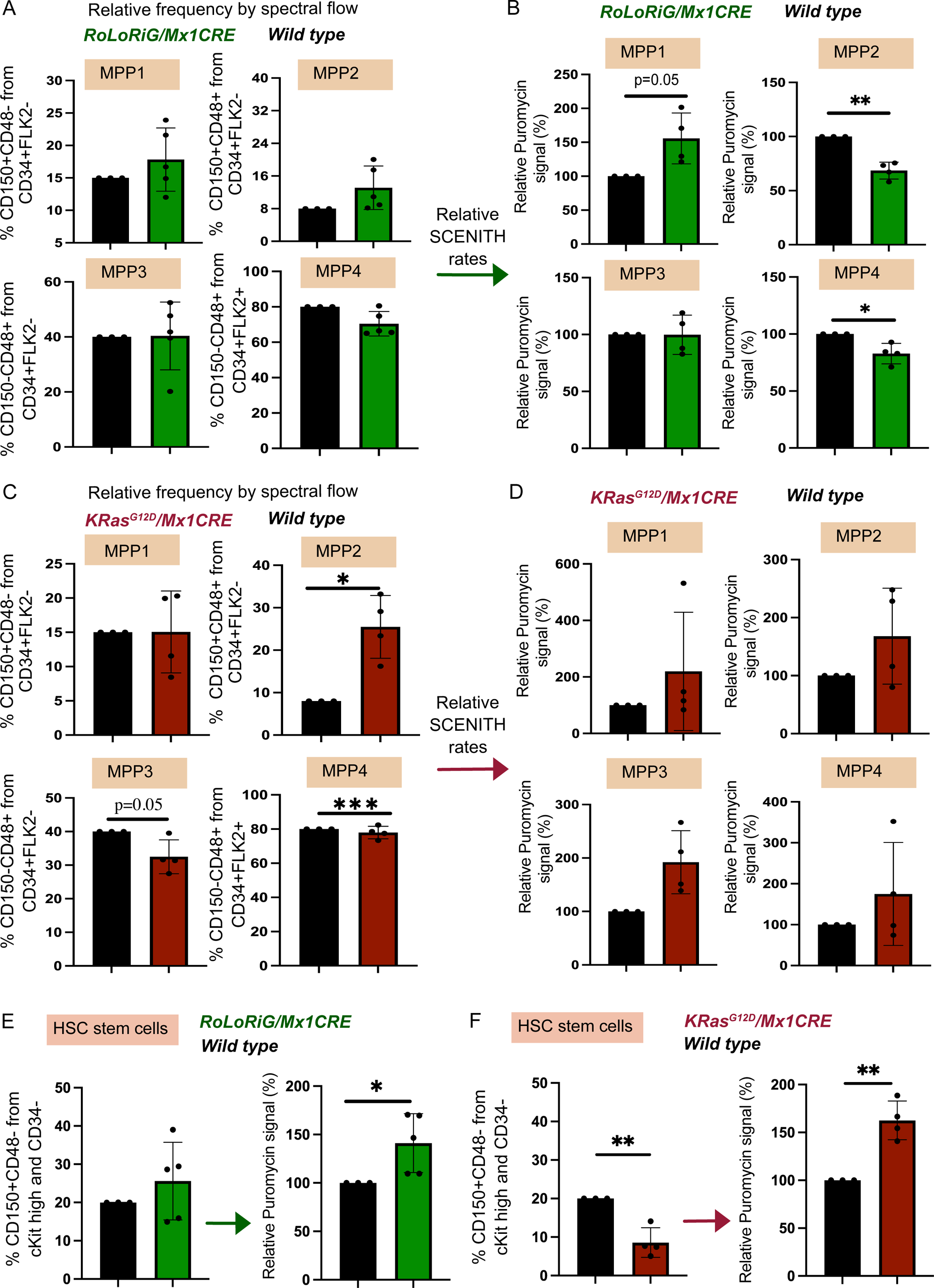
Impact of Ras pathway lesions on MPP- and HSC stem cells and their SCENITH™ signals. (A) Relative frequency of MPP1-4 progenitor populations for RoLoRiG/Mx1CRE and wild type mice. Wild type control MPP frequency were arbitrarily set at 15%, 8%, 40%, and 808%, respectively. See Supplemental Figure S12. (B) Relative SCENITH™ signals in RoLoRiG/Mx1CRE and wild type MPP1-4 progenitor populations. Mean fluorescence Puro signals were determined in the MPP populations defined in 7A, wild type MPP SCENITH™ signals were arbitrarily set at 100% in each MPP population. *p<0.05; **p<0.01. (C) As in 7A but for MPP1-4 progenitor populations from KRASG12D/Mx1CRE and wild type mice. *p<0.05. (D) Reltive SCENITH™ signals in MPP1-4 progenitors from KRASG12D/Mx1CRE and wild type mice. As in 7B. (E) Relative frequency of HSC stem cells and relative SCENITH™ signals in these HSC for RoLoRiG/Mx1CRE and wild type mice. Wild type control HSC frequency was arbitrarily set at 20%, see Supplemental figure S10E and F, with a relative SCENITH™ signal of 100%. *p<0.05. (F) As in 7E but comparing KRASG12D/Mx1CRE (n = 3) HSC and against 3 other wild type HSC. Note that both the fraction of KRASG12D/Mx1CRE HSC in the LSK compartment and the LSK fraction itself were reduced in KRASG12D/Mx1CRE mice, see Supplemental Figure S10E and F. **p<0.01.

Finally, for HSC, defined here as a CD150-positive/CD48-negative fraction from the cKit-positive/CD34-negative LSK population (**Supplemental Figure S10E**), the *RoLoRiG/Mx1CRE* BM revealed a trend of an increased fraction of HSC at four to eight weeks post pIpC injection (**Supplemental Figure S10F**) and these HSC displayed increased protein synthesis levels (**Figure 7E**). By contrast, not only is the *KRAS^G12D^/Mx1CRE* LSK compartment diminished (**Figures 2H**, **3C, Supplemental Figure S8E,** and **6E**), the HSC population with the *KRAS^G12D^* lesion - calculated as a fraction from LSK- is further reduced and *KRAS^G12D^/Mx1CRE* HSC displayed increased protein synthesis levels (**Figure 7F**).

## DISCUSSION

As mentioned, Ras pathway lesions are highly prevalent in hematopoietic malignancies (Bowen et al., 2005; Liu et al., 2017; Schubbert et al., 2007; von Lintig et al., 2000; Ward et al., 2012). Leukemia cell lines expressing KRas^G12D^ display high, constitutive levels of active RasGTP, whereas leukemia lines overexpressing RasGRP1 do not but, instead, display robust RasGTP levels when receiving cytokine receptor signals (Hartzell et al., 2013). Mouse models of oncogenic Ras mutation in the bone marrow (Bowen et al., 2005; Braun et al., 2004; Chan et al., 2004; Kindler et al., 2008; Li et al., 2011; Liu et al., 2017; Sabnis et al., 2009; Schubbert et al., 2007; Wang et al., 2010), like *KRas^G12D^* and our new *RoLoRiG/Mx1CRE* strain (Karra et al., 2020) have provided a framework of aberrant Ras signals in the hematopoietic compartment, but the precise impacts of such Ras pathway lesions on rare HSC and LSK cells are not known.

Here we provided single cell proteomics and metabolomics datasets as a resource of mechanistic insights into altered hematopoiesis driven by single, distinct, induced Ras pathway lesions. In normal hematopoiesis, dormant HSC enter the cell cycle upon tissue insults, such as exposure to chemotherapy 5-FU (5-Fluoro-Uracil) or bone marrow transplantation, but re-enter dormancy after the blood system is repaired (Wilson et al., 2008). For these reasons we performed our studies in the context of native hematopoiesis, i.e., without transplantation, capitalizing on *KRAS^G12D^/Mx1CRE*- and *RoLoRiG/Mx1CRE*-mice and pIpC injection.

At first glance, the bone marrow of *KRAS^G12D^/Mx1CRE*- and *RoLoRiG/Mx1CRE*-mice appears very similarly altered since both models show a robust skewing towards CD11b-positive myeloid cells, albeit that the KRas^G12D^-driven myeloid expansion is aggressive and overexpression of RasGRP1 has a milder effect. The resolution of CyTOF allowed for detailed examination of very rare Stem- and progenitor- cells, and we established the surprising opposing phenotypes of an expanded LSK- and HSC-compartment in *RoLoRiG/Mx1CRE* mice and a depletion of these compartments in *KRAS^G12D^/Mx1CRE* mice. These high-resolution characterizations are in agreement with earlier, lower-resolution reports that *KRas^G12D^* induction results in a decrease in LSK numbers (Sabnis et al., 2009) and that overexpression of RasGRP1 bestows some sort of fitness to stem cells (Karra et al., 2020).

It should be noted that pIpC injection of *NRas^G12D/WT^/Mx1CRE* mice has been reported to yield a very mild, delayed myeloproliferative neoplasm (Li et al., 2011; Wang et al., 2013; Wang et al., 2011; Ward et al., 2012) with similarities to *RoLoRiG/Mx1CRE*-mice. Interestingly, *NRas^G12D/G12D^/Mx1CRE* mice with two mutated alleles consistently develop a similar MPD as *Kras^G12D/+/^MX1CRE* mice (Xu et al., 2013; Zambetti et al., 2020). Similarly, knock-in mice expressing the lesser activating *KRas* mutations A146T (Poulin et al., 2019) and T58I (Wong et al., 2020) retain near-normal numbers of HSC. Together with our data, these studies support the idea that the degree and duration of Ras signal output has profound effects on stem cell maintenance and lineage-potential.

SCENITH™ analysis provided mechanistic insights into the rare, lineage-negative immature cells and deeper metabolic characterization of LSK and HSC compartments. In both the *KRAS^G12D^/Mx1CRE*- and *RoLoRiG/Mx1CRE*-mice, we see an elevated level of protein synthesis in lineage-negative compartments and in HSC stem cells, compared to *wild type* controls. Different between the two models with distinct Ras pathway lesions is the depletion of metabolically quiescent Stem- and progenitor- cells when KRas^G12D^ is present but the maintenance of that quiescent population in *RoLoRiG/Mx1CRE* mice. These metabolic characterizations with single cell resolution offer mechanistic explanations for several reported but unexplained observations. First, in *RoLoRiG/Mx1CRE* mice, RasGRP1-overexpressing cells (that also express GFP) take over the entire hematopoietic production over time and, thus, somehow have an advantage over GFP-negative cells (Karra et al., 2020). Second, *KRas^G12D^* LSK cells exhibit a 10-fold reduction in long-term repopulation capacity (Sabnis et al., 2009) and transplantation of *KRas^G12D^/Mx1CRE* cells into sub-lethally irradiated recipient mice do not reconstitute hematopoiesis (Braun et al., 2004). Of note, *NRas^G12D^/Mx1CRE* HSC reveal a bimodal cycling pattern (Li et al., 2013) and outcompete *wild type* cells in transplantation assays (Wang et al., 2013; Wang et al., 2011).

The strong, constitutive KRas^G12D^ signal results not only in an elevated, mean level of protein synthesis but also in loss of the SCENITH™-low state. Possibly, driving leukemic cells out of SCENITH™-low state can be explored as avenue for future therapies. The distinct SCENITH™ patterns of LSK and HSC compartments in *KRAS^G12D^/Mx1CRE*- and *RoLoRiG/Mx1CRE*-mice also imply that successful leukemias in patients with either KRas^G12D^ or RasGRP1 overexpression must pair with distinct sets of complementary genetic lesions that balance the protein synthesis rates observed with distinct Ras pathway lesions. Indeed, mouse model leukemia virus insertion studies demonstrated that KRas^G12D^ or RasGRP1 overexpression desire distinct insertions in other gene loci (Dail et al., 2010; Hartzell et al., 2013; Karra et al., 2020; Lauchle et al., 2009). It will be of interest to investigate how these other genes impact the metabolism of Stem- and progenitor- cells in the bone marrow.

## ACKNOWLEDGMENTS

This work was supported by following awards: American Association for Cancer Research award 20-20-01-SPIT, Cancer Research Institute award CRI4437, NIH award R01DE032033, American Cancer Society award RSG-22-141-01-IBCD, DOD US Army Med. Res. Acq. Activity Award BC220499, and NIH award DP5 OD023056 (all to MHS). The CyTOF instrument used in this study was purchased with assistance by NIH award S10 1S10OD018040-01. This work has been supported by CENTURI postdoc funding PGG. Also, work was supported by the ANR JCJC-Epic SCENITH N° ANR-20-CE14-0028 to RJA; ANR PRC MetaNiche N° ANR-22-CE15-0015-02 to RJA and European Commission Horizon2020 Transcan2021-227-TALETE French National Cancer Institute (INCa) to RJA. Further, this project was conceptualized with support work from an Alex’ Lemonade Stand Foundation Innovator Award with main funds provided by NIH/NCI (R01 - CA187318), and support funds from a NIH/NIAID (P01 AI091580) (all to JPR). Further support for the Roose lab came from a Rothschild Fellowship for postdoctoral fellows in the Natural, Exact or Life Sciences and Engineering (to LK) and a Momentum Fellowship from the Mark Foundation for Cancer Research (to A-MF). We thank the members of the Combes-, Arguello-, Spitzer-, and Roose-labs for helpful input. We thank our UCSF colleagues Drs. Serine Avagyan and Kevin Shannon for proofreading the manuscript.

## AUTHOR CONTRIBUTIONS

LK collected BM samples, performed experiments, analyzed results, and wrote draft sections of the manuscript. A-MF, LS, and MK performed SCENITH™ experiments and analyzed and plotted results. RMY analyzed CyTOF data. MP, KB, LH assisted with various Spectral Flow experiments and mouse breeding. IT conjugated and titrated antibodies for CyTOF and generated CyTOF data. RJA and PGG produced and tested the reagents and provided key assistance with the SCENITH™ methodology, data interpretation and discussion on SCENITH and data. MS designed and supervised CyTOF data generation and analysis. AJC provided advice on SCENITH™ experiments and HG on general experimental approaches. LK, A-MF, MK, RMY generated figure panels. JPR designed the research program, secured most of the funding, and wrote the final draft manuscript. LK, A-MF, LS, AJC, HG, RJA, MHS provided manuscript edits.

## Competing Interests Statement

MHS is founder, shareholder and board member of Teiko.bio, has received a speaking honorarium from Fluidigm Inc., Kumquat Bio, and Arsenal Bio, has been a paid consultant for Five Prime, Ono, January, Earli, Astellas, and Indaptus, and has received research funding from Roche/Genentech, Pfizer, Valitor, and Bristol Myers Squibb. JPR is a co-founder and scientific advisor of Seal Biosciences, Inc., a scientific advisory committee member for the Mark Foundation for Cancer Research, a consultant for MorphImmune and Monte-Rosa Therapeutics, and has received research funding from 3TBiosciences, Senti Biosciences, and Eli Lilly.

## SUPPLEMENTARY MATERIAL

- Four Supplementary Tables
- Resource Table
- Ten Supplemental Figures

## SUPPLEMENTAL FIGURE LEGENDS

**Figure S1: Single cell proteomics characterization of total bone marrow of *wild type, RoLoRiG/Mx1CRE,* and *KRAS^G12D^/Mx1CRE* mice using Phenograph analysis.**

(A) Schematic representation of altered Ras signals.

KRas^G12D^ is severely impaired in its self-inactivating GTPase function.

(B) Schematic of hematopoiesis in the bone marrow (BM).

(C) 40 surface markers used in our CyTOF approach.

(D) Schematic explanation of collapsing 40 dimensions into UMAPs.

(E, F) UMAP representation of cell composition in *wild type* BM, *RoLoRiG/Mx1CRE* total BM at 4 weeks post-pIpC injection, and *KRAS^G12D^/Mx1CRE* total BM at 4 weeks post-pIpC injection. UMAPs are compilations of n = 4 mice.

**Figure S2: PhenoGraph analysis of total BM cell populations.**

(A) PhenoGraph analysis of *Wild type* total BM. Color-coded clusters are generated based on protein expression values, revealing 26 clusters with assigned cell compositions. Figure S2A depicts the total BM composite of four *wild type* mice (See **Supplemental Figure S3** for PhenoGraphs of all twenty individual mice).

(B) PhenoGraph maps of *RoLoRiG/Mx1CRE* total BM (4 weeks or 4 months after pIpC, compilation of n = 4 mice each).

(C) PhenoGraph maps of *KRAS^G12D^/Mx1CRE* total BM (4 weeks or 2 months after pIpC, compilation of n = 4 mice each). In both *RoLoRiG/Mx1CRE* and *KRAS^G12D^/Mx1CRE* BM, B cells and neutrophils drastically decrease at the late post-pIpC time points (clusters #1, 2, and 9).

(D) Aitchitson analysis depicting the level of compositional divergence of *RoLoRiG/Mx1CRE* and *KRAS^G12D^/Mx1CRE* total BM, compared to total *wild type* BM. *p<0.05.

**Figure S3: PhenoGraph analysis of total BM cell populations in twenty individual mice.**

PhenoGraph analysis of total BM cell populations in twenty individual, *Wild type* (#1-4) *RoLoRiG/Mx1CRE* (#5-8 and #9-12), or *KRAS^G12D^/Mx1CRE* (# 13-16 and #17-20) mice. As in Supplemental Figure S2, samples are clustered based on protein expression values, revealing 26 clusters with assigned cell compositions. *RoLoRiG/Mx1CRE* mice were analyzed 4 weeks or 4 months post-pIpC injection. *KRAS^G12D^/Mx1CRE* mice were analyzed 4 weeks or 2 months post-pIpC injection. Color-coding for cell subsets as in Supplemental Figure S2.

**Figure S4: Hierarchical clustering of marker expression in 26 cell subset clusters.**

(A) Examples of individual marker expression intensity overlayed onto Phenographs.

(B) Extraction of quantitative maker values and conversion into heatmaps (for details see Supplemental Figure S5)

(C) Hierarchical clustering of 26 cell subsets (in rows) against the 40 markers and their values (in columns). Depicted here is *wild type* total bone marrow, for comparisons to total BM with Ras pathway lesions, see Supplemental Figure S5.

**Figure S5: Hierarchical clustering and heatmap presentation of 26 cell subset clusters and 40 markers.**

Using the PhenoGraph analysis that revealed 26 clusters with assigned cell compositions, heat maps of the resulting cluster frequencies and hierarchical clustering of the heat maps were generated with the Seaborn package in Python. Log2 fold change heat maps were calculated as the log2 of the fold change in cluster frequencies between the indicated two groups.

(A) Hierarchical clustering of heat maps for total BM cell populations in *wild type* mice.

(B, C) Hierarchical clustering of heat maps for total BM cell populations in *RoLoRiG/Mx1CRE* mice, analyzed 4 weeks (B) or 4 months (C) post-pIpC injection.

(D, E) Hierarchical clustering of heat maps for total BM cell populations in *KRAS^G12D^/Mx1CRE* mice, analyzed 4 weeks (D) or 2 months (E) post-pIpC injection.

(F) Marker composition of clusters #3, #15, #16, #22, #23, #25 in total BM, corresponding to Figures 1H. Clusters #22, 23, and 25 have unique properties but share unusual co-expression of T-cell (TCRb, CD3, CD4 and CD8) and B cell (IgM, IgD, B220) markers. Clusters #22, 23, 25 were very rare in *wild type* mice, but increased at 4 months for the *RoLoRiG/Mx1CRE* mice and at both 4 weeks and 2 months in *KRAS^G12D^/Mx1CRE mice* (See Supplemental Figure S2). We termed these clusters “DEs or dual expressers”, following the term used in studies of type-1 diabetic patients (Ahmed et al., 2021). The existence of cluster #3 is identified by PhenoGraph but datapoints are negligible. Throughout this study we also observe several cell subsets, such as clusters #15 and #16 in total BM, that we cannot classify with confidence; we term these “undefined” but list the complete marker expression profiles as an unbiased resource.

**Figure S6: Comparisons of cell subset cluster frequencies in total BM.**

(A) To comprehensively portray the overall changes in total BM, we plotted the relative abundance of the 26 cell clusters, visualizing overall changes in the total bone marrow compartment. For example, note the reduction in B cells (cluster #2 in blue) in total *RoLoRiG/Mx1CRE* and total *KRAS^G12D^/Mx1CRE* BM, analyzed at the four time points.

(B) Color-coded visualization of Log2 fold changes in cell cluster frequencies in total *RoLoRiG/Mx1CRE* and *KRAS^G12D^/Mx1CRE* BM, compared to *Wild type*. Note the similar pattern of decreases (in blue) and increases (in red) for the two Ras pathway lesions, compared to *wild type*.

(C) Head-to-head comparisons of cell cluster frequencies, analyzing the differences between *RoLoRiG/Mx1CRE* and *KRAS^G12D^/Mx1CRE* at the indicated time points post-pIpC injection.

**Figure S7: PhenoGraph-assisted profiling of the Myeloid BM compartment.**

(A) Same image as Figure 1H. PhenoGraph analysis revealing 23 CD11b-positive clusters in BM of *wild type* mice, compilation of n = 4 mice. Cluster #10 and 11 are identified by PhenoGraph but datapoints are negligible and #5 is undefined (See **Supplemental Table S2**). (B-C) As in S6A, but for *RoLoRiG/Mx1CRE* and *KRAS^G12D^/Mx1CRE*, with the indicated periods post-pIpC injection. Each figure is a compilation of n = 4 mice. Arrows point out cluster #1 neutrophils.

(D) Color-coded visualization of Log2 fold changes in cell cluster frequencies in CD11b-positive *RoLoRiG/Mx1CRE* and *KRAS^G12D^/Mx1CRE* BM, compared to *Wild type*. Note the similar pattern of decreases (in blue) and increases (in red) for the two Ras pathway lesions, compared to *wild type*.

**Figure S8: Analyses of the lineage-negative compartment in the bone marrow.**

(A) Schematic representation of gating strategy of CyTOF data to analyze lineage-negative cells in the BM.

(B) PhenoGraphs of lineage-negative cells showing 20 cell clusters in BM from *RoLoRiG/Mx1CRE* or *KRAS^G12D^/Mx1CRE* mice, 4 weeks after pIpC injection. Each figure is a compilation of n = 4 mice. The respective increases in CMP in the *RoLoRiG/Mx1CRE* and *KRas^G12D^/Mx1CRE* models closely mirrored the magnitude of elevated CD11b-positive cell fractions (**Figure 1K**), suggesting that the aberrantly expanded CMP generally follow maturation to CD11b-positive myeloid cell lineages in the bone marrow of both models. A notable exception was #2 MEP (Megakaryocyte/erythroid progenitors) that give rise to platelets and erythrocytes. MEP are born out of CMP but did not mirror the CMP pattern in *RoLoRiG/Mx1CRE* and *KRas^G12D^/Mx1CRE* models. In *KRAS^G12D^/Mx1CRE mice*, MEPs decreased significantly at both 4 weeks and 2 months. MEP were increased 4 weeks post-pIpC but decreased at 4 months in *RoLoRiG/Mx1CRE* mice (**Supplemental Figure S8B and Figure 2F**).

(C) Compositional Aitchinson distances in cell populations of the lineage-negative compartment in the different mouse models.

(D) Murine hematopoiesis: representation of the LSK (Lineage negative, Sca1 and cKit positive) compartment, depicting the updated gating strategy on the left and the more traditional gating strategy on the right. In this study, we used the cell subset analysis in which MPPs 1-4 arise from HSCs and give rise to more mature progenitors with restricted cell fate, as published in (Wilson et al., 2008).

(E) PhenoGraph analysis of the LSK compartment for *KRAS^G12D^/Mx1CRE* BM, 4 weeks post-pIpC injection.

(F) Representation of the approximate fraction (%) of cells from subsequent gates to illustrate the scarcity of LSK and HSC cells. Note that the frequency of the numbers here was based on our CyTOF data from *wild type* mice.

**Figure S9: Analyses of the LSK compartment in the bone marrow by CyTOF.**

(A) Compositional Aitchinson distances in cell populations of the LSK compartment in the different mouse models.

(B) Frequencies of MPP1, CLP + CLP-like, MPP3-like, MPP4-like, #10 Undefined; CD90+ cells, and #7 Undefined CD44-cells as a percentage of LSK cells for the indicated experimental groups. *p<0.05 **p<0.01 ***p<0.001 ****p<0.0001.

(C) Schematic of LSK cell subsets that we can detect, discriminate, and quantitate with our CyTOF, not depicting #10 Undefined; CD90+ cells and #7 Undefined CD44-cells.

(D-F) Relative percentages of specific cell subsets within the LSK population for the three genetic mouse models with significant alterations in colors with statistical analysis. *p<0.05 **p<0.01 ***p<0.001 ****p<0.0001.

**Figure S10: SCENITH™ in bone marrow and unmixing and compensation of BM spectral flow**

(A) Representative SCENITH™ histograms of *RoLoRiG/Mx1CRE* and *wild type* total bone marrow. Gates were selected using the positive spectral flow SCENITH™ values. A Puromycin-low gate was set, starting at the value zero and moving rightwards until 20% of *wild type* cells fell in that gate. Vice versa, a Puromycin-high gate was set, starting at the highest SCENITH™ value and moving leftwards until 20% of *wild type* cells fell in that gate. Gates were copied over to *RoLoRiG/Mx1CRE* total bone marrow that was run in parallel on the same day. These gates or fractions of Puromycin-low and Puromycin-high total bone marrow cells were arbitrarily set at 0.2 for wild type and used for Figure 5D.

(B) Same as in S10A, but for *KRAS^G12D^/Mx1CRE* and *wild type* total bone marrow to calculate values for Figure 5E.

(C) Experimental pipeline and data analysis approach to unmix, compensate, and visualize SCENITH™ analysis in defined and rare cell subsets through the combination with Spectral flow.

(D) Process of Spectral Flow data analysis, see our previous work (Hysenaj et al., 2023).

(E) Gating strategy for our BM Spectral Flow stainings. Following exclusion of lineage-positive, mature cells the roughly 35% of lineage-negative, immature cells are further sub-setted into LSK Stem- and progenitor- cells and into HSC stem cells.

(F) Relative frequencies on HSC cell numbers, calculated as CD150-positive/CD48-negative cells as a fraction of the cKit-high/CD34-negative cells. For each pair of mice in every experiment, *wild type* HSC were arbitrarily set at 20% and the corresponding percentage was calculated for the models with Ras pathway lesions. These relative HSC fractions were plotted in Figures 7E and 7F.

**Figure S11: t-SNE plots for SCENITH™ analyses of the lineage-negative BM compartment with Ras pathway lesions.**

As in Figure 5F-H and 5I-K.

(A-B) t-SNE plots representing cell composition, CD34 expression levels, and SCENITH™ levels for lineage negative BM cells from individual *RoLoRiG/Mx1CRE* and *wild type* mice. Note that (i) individual *RoLoRiG/Mx1CRE* and *wild type* mice are always compared in experimental pairs, (ii) that every experiment of a pair of test- and control-mouse is an individual experiment with a unique puromycin incorporation characteristic (this is why we always compare pairs), and (iii) that the anti-Puromycin antibody here is coupled to Alexa647. (C-D) As in S11A-B, but for the *KRAS^G12D^/Mx1CRE* and *wild type* lineage-negative compartment. Note that the anti-Puromycin antibody here is coupled to Alexa488.

**Figure S12: Spectral flow analysis of MPP1, MPP2, MPP3, and MPP4 populations.**

Gating strategy for MPP1, MPP2, MPP3, and MPP4 populations with our BM Spectral Flow stainings, based on the indicated markers. Calculation of relative frequency of MPP1-4 progenitor populations in the indicated mouse models. *Wild type* control MPP frequency were arbitrarily set at 15%, 8%, 40%, and 808%, respectively. MPP populations in the test mice with Ras pathway lesions were determined from gates that were identically copied over from *wild type* control mice.

