## Supplemental Figures and Tables for "Single cell proteomics characterization of bone marrow hematopoiesis with distinct Ras pathway lesions"

A

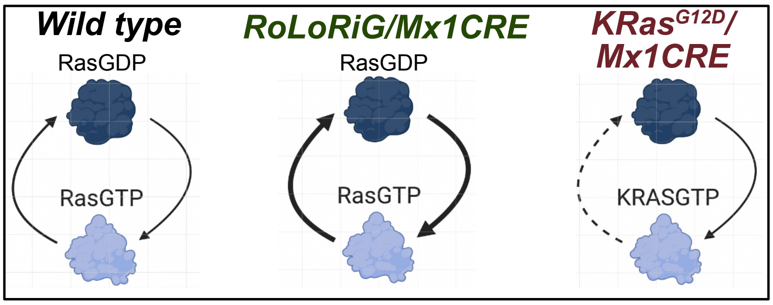

B

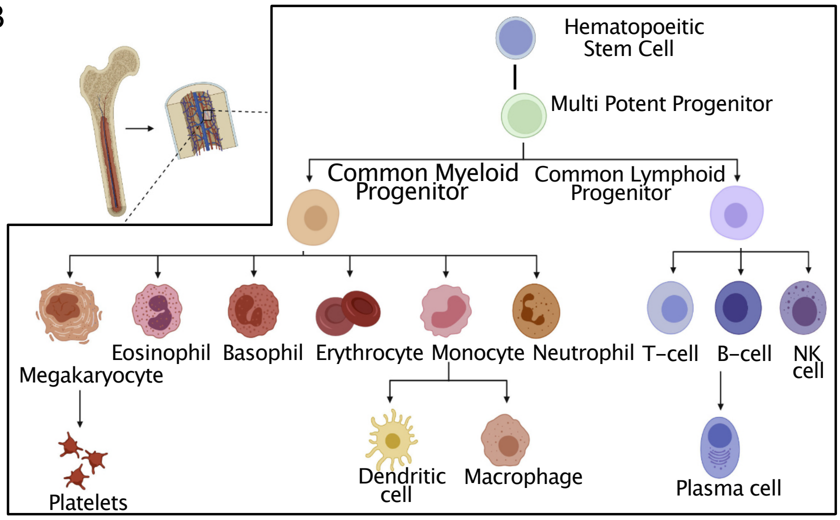

C

| Surface Markers | | | |
|---|---|---|---|
| *CD45.1* | CD41 | TCRb | F4/80 |
| *CD45.2* | SiglecF | B220 | CD115 |
| Ly6G | CD90 | CD127 | CD71 |
| IgD | PDCA-1 | *Ter119* | CD23 |
| CD16/32 | Ly6C | MHC II | CD19 |
| CD49b | CD48 | CD105 | IgM |
| CD11c | CD11b | Sca1 | CD44 |
| CD43 | cKit | CD150 | |
| CD27 | CD8 | FcER1a | |
| CD138 | CD4 | CD135 | |
| CD34 | CD3 | CD25 | |

CyTOF

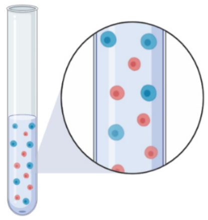

+ 20 Mass-tag barcodes

D

CyTOF proteomics datasets with 40 dimensions

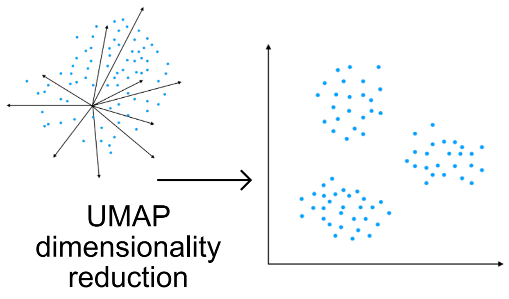

E

Compilation of n = 4 mice

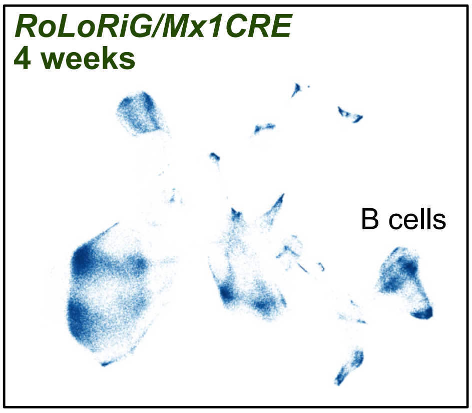

F

Compilation of n = 4 mice

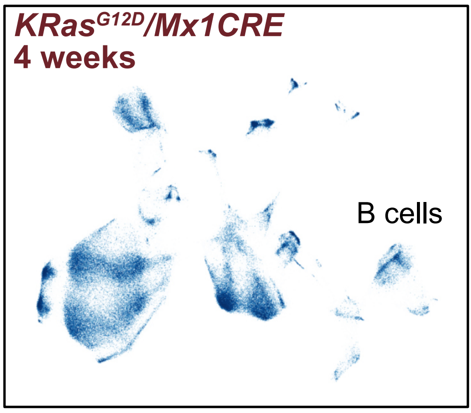

**Karra *et al.* , Supplemental Figure S1**

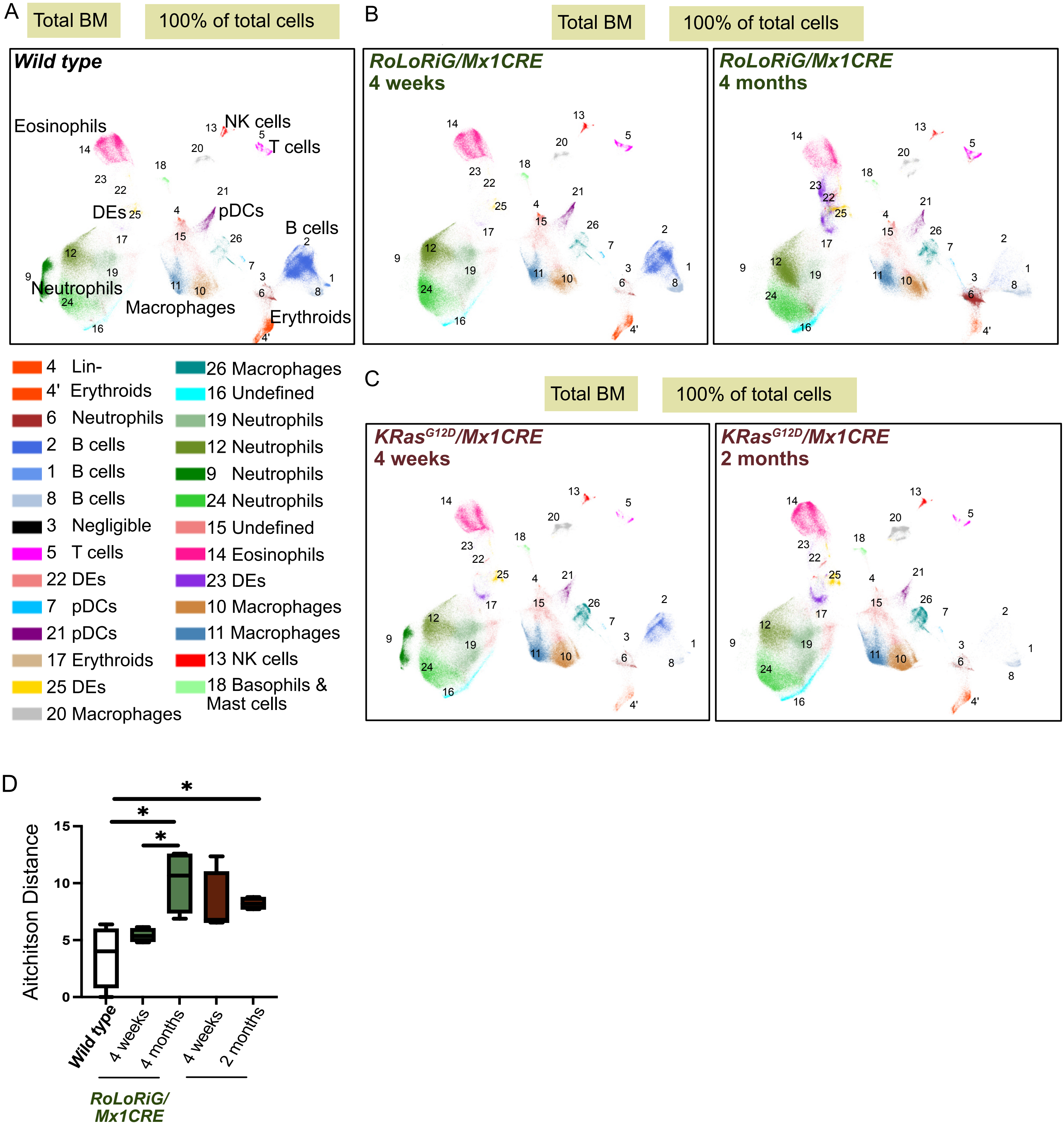

**Karra *et al.* , Supplemental Figure S2**

Phenographs of individual mice

**Total BM** **100% of total cells**

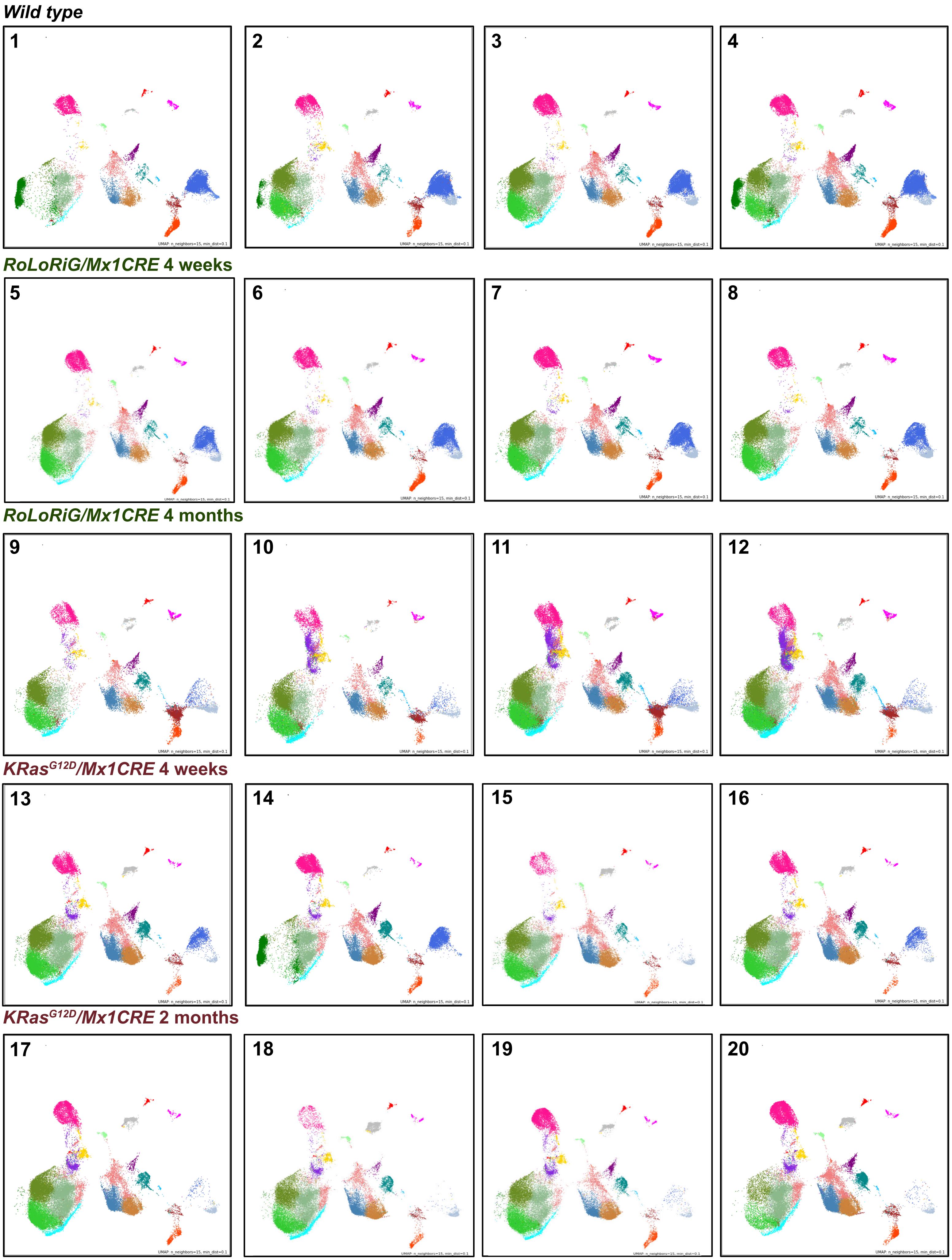

**Karra *et al.* , Supplemental Figure S3**

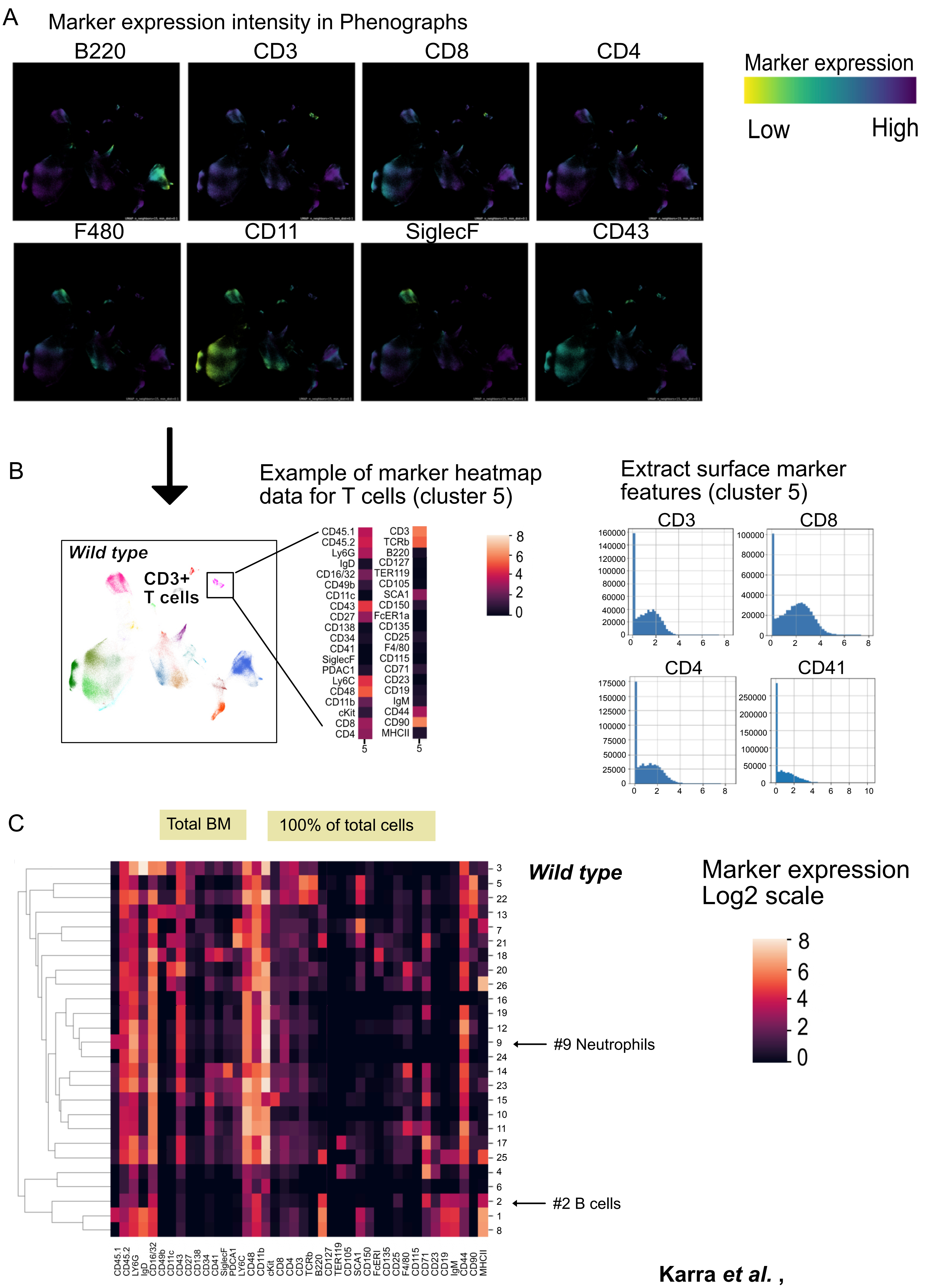

**Karra *et al.* ,**
**Supplemental Figure S4**

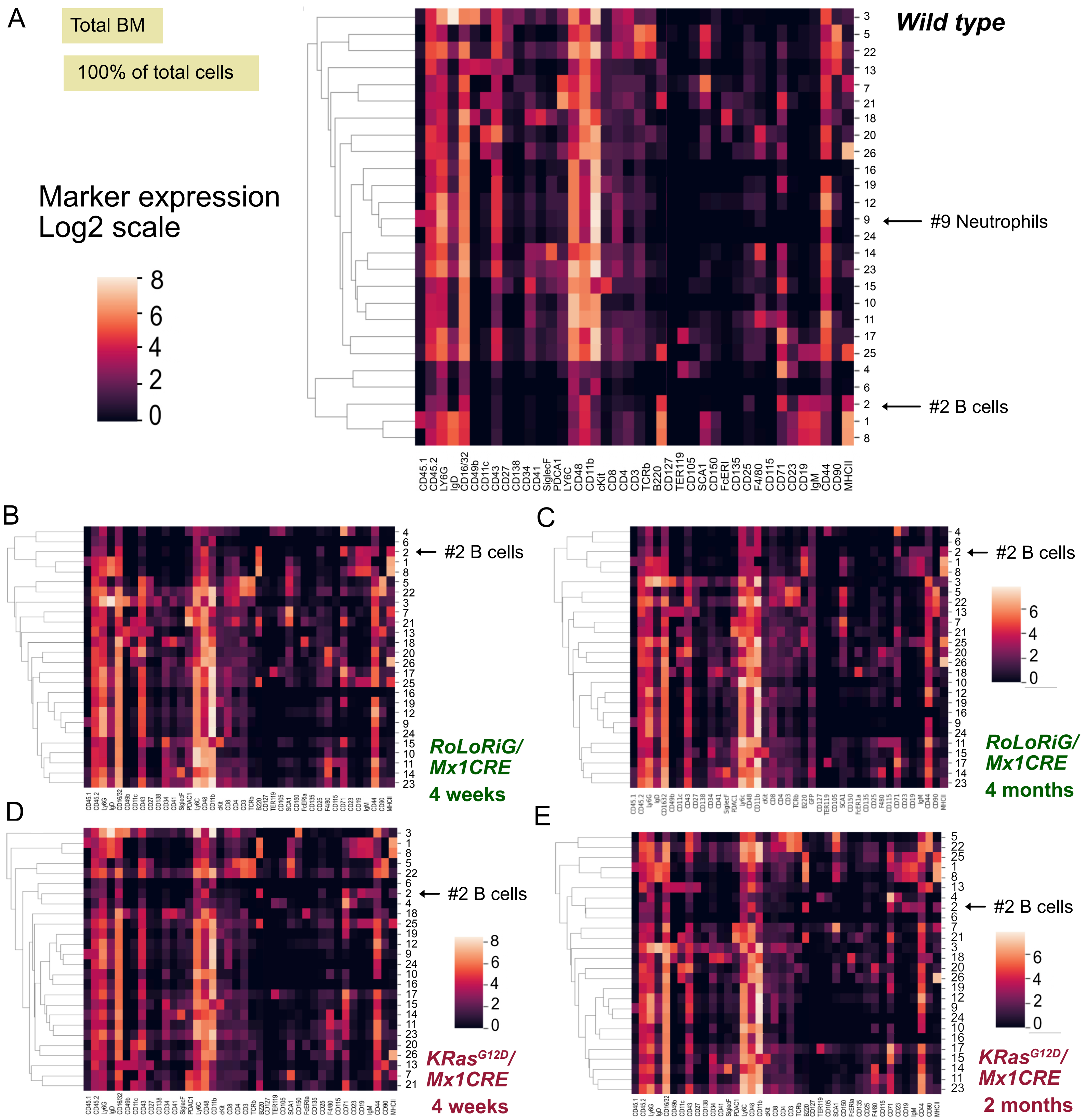

F

| Cluster number | Markers expressed |
|---|---|
| 3 | MHCII$^{low}$ CD44$^{+}$ IgM$^{low}$ CD150$^{+}$ TCRb$^{low}$ CD4$^{+}$ CD8$^{+}$ ckit$^{low}$ CD11b$^{+}$ CD48$^{med}$ Ly6C$^{+}$ PDCA1$^{low}$ CD41$^{low}$ CD34$^{low}$ CD138$^{low}$ CD27$^{low}$ CD43$^{+}$ CD11c$^{low}$ CD49b$^{+}$ CD16/32$^{+}$ IgD$^{+}$ Ly6G$^{+}$ |
| 15 | CD44$^{med}$ CD71$^{med}$ F480$^{low}$ CD25$^{low}$ SCA1$^{low}$ CD105$^{low}$ CD3$^{low}$ CD4$^{low}$ CD8$^{low}$ cKit$^{+}$ CD11b$^{low}$ CD48$^{+}$ Ly6C$^{+}$ PDCA1$^{low}$ SiglecF$^{low}$ CD41$^{low}$ CD34$^{+}$ CD27$^{low}$ CD43$^{med}$ CD16/32$^{+}$ Ly6G$^{med}$ |
| 16 | CD3$^{low}$ CD8$^{low}$ cKit$^{low}$ CD11B$^{+}$ CD48$^{med}$ Ly6C$^{+}$ SiglecF$^{low}$ CD34$^{low}$ CD43$^{+}$ CD16/32$^{+}$ IgD$^{low}$ Ly6G$^{+}$ |
| 22 | MHCII$^{low}$ CD90$^{+}$ CD44$^{+}$ IgM$^{low}$ CD71$^{low}$ F480$^{low}$ CD25$^{low}$ CD150$^{low}$ SCA1$^{med}$ B220$^{low}$ TCRb$^{+}$ CD3$^{+}$ CD4$^{med}$ CD8$^{med}$ cKit$^{low}$ CD11b$^{+}$ CD48$^{+}$ Ly6C$^{+}$ PDCA1$^{low}$ SiglecF$^{low}$ CD41$^{low}$ CD34$^{low}$ CD27$^{low}$ CD43$^{+}$ CD11C$^{low}$ CD49b$^{low}$ CD16/32$^{+}$ IgD$^{low}$ Ly6G$^{+}$ |
| 23 | MHCII$^{low}$ CD90$^{low}$ CD44$^{+}$ CD71$^{low}$ F4/80$^{med}$ CD25$^{low}$ SCA1$^{low}$ B220$^{low}$ TCRb$^{low}$ CD3$^{low}$ CD4$^{low}$ CD8$^{med}$ cKit$^{low}$ CD11b$^{+}$ CD48$^{+}$ Ly6C$^{+}$ PDCA1$^{med}$ SiglecF$^{low}$ CD41$^{low}$ CD34$^{med}$ CD27$^{low}$ CD43$^{+}$ CD16/32$^{+}$ IgD$^{low}$ Ly6G$^{+}$ |
| 25 | MHCII$^{+}$ CD44$^{+}$ IgM$^{med}$ CD19$^{med}$ CD71$^{+}$ F4/80$^{low}$ CD25$^{low}$ SCA1$^{low}$ B220$^{+}$ CD3$^{low}$ CD4$^{low}$ CD8$^{med}$ cKit$^{low}$ CD11b$^{+}$ CD48$^{+}$ Ly6C$^{+}$ PDCA1$^{low}$ SiglecF$^{low}$ CD34$^{low}$ CD43$^{+}$ CD16/32$^{+}$ IgD$^{low}$ Ly6G$^{+}$ |

**Karra *et al.* , Supplemental Figure S5**

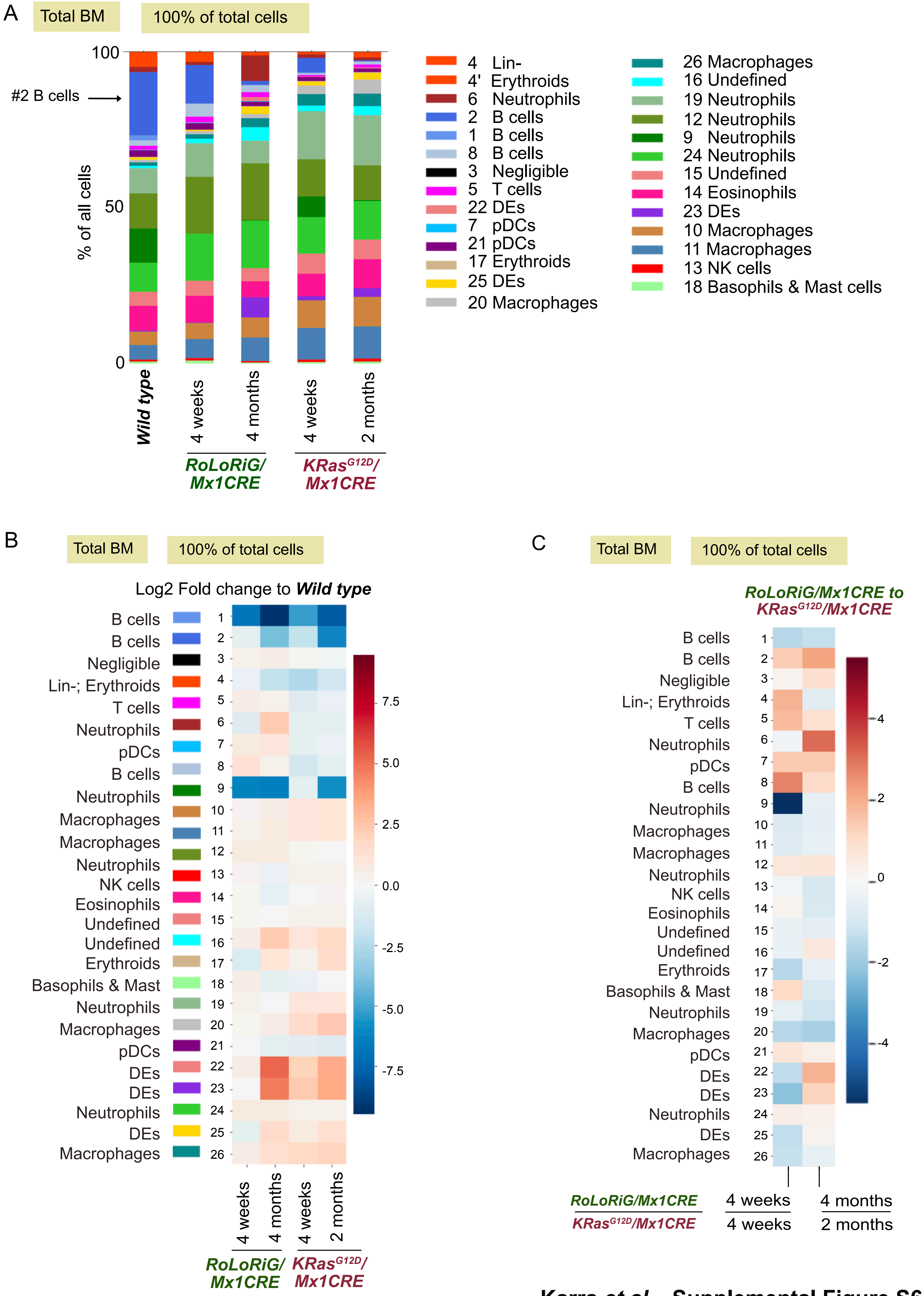

**Karra *et al.* , Supplemental Figure S6**

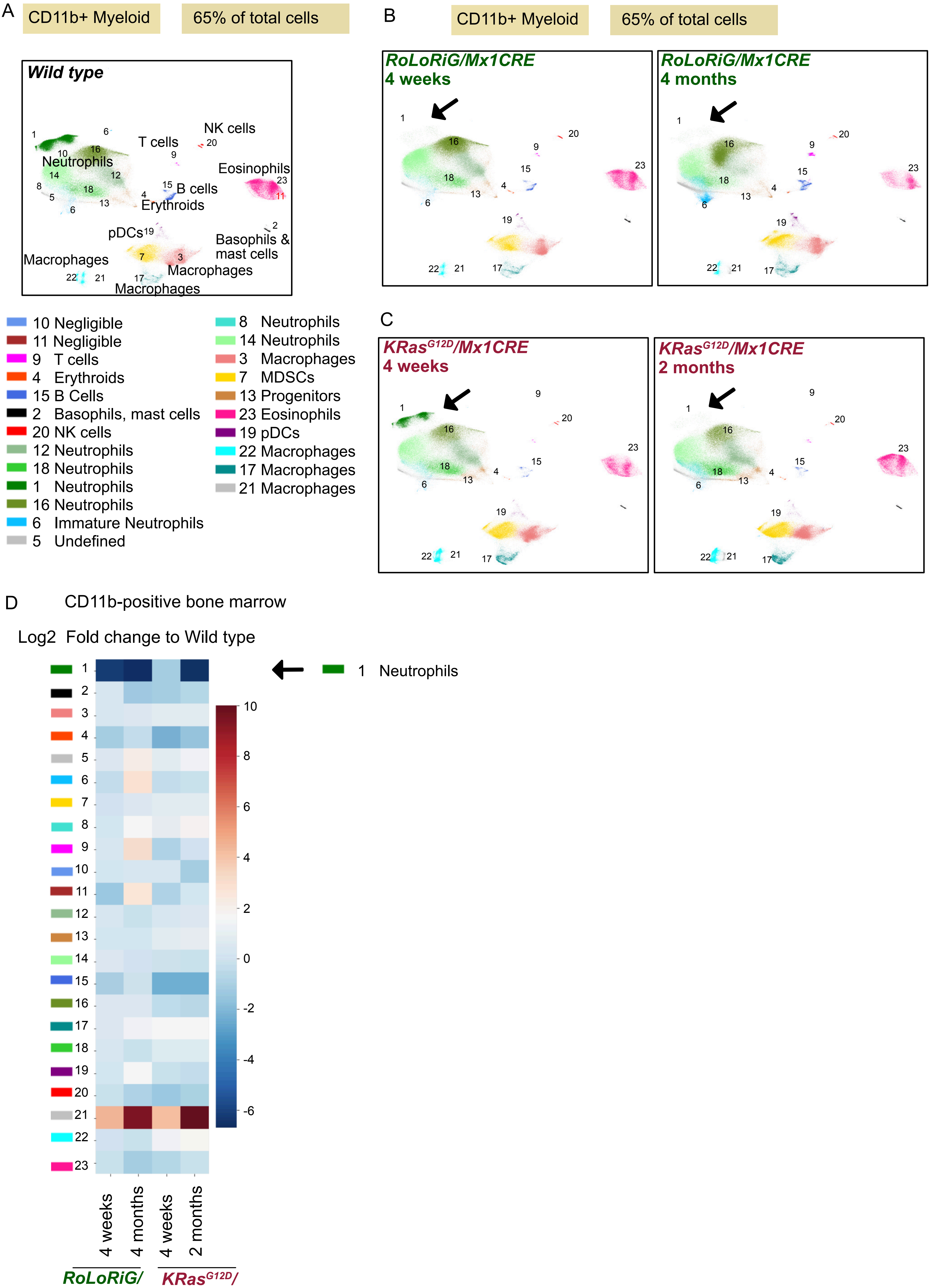

Karra *et al.* , Supplemental Figure S7

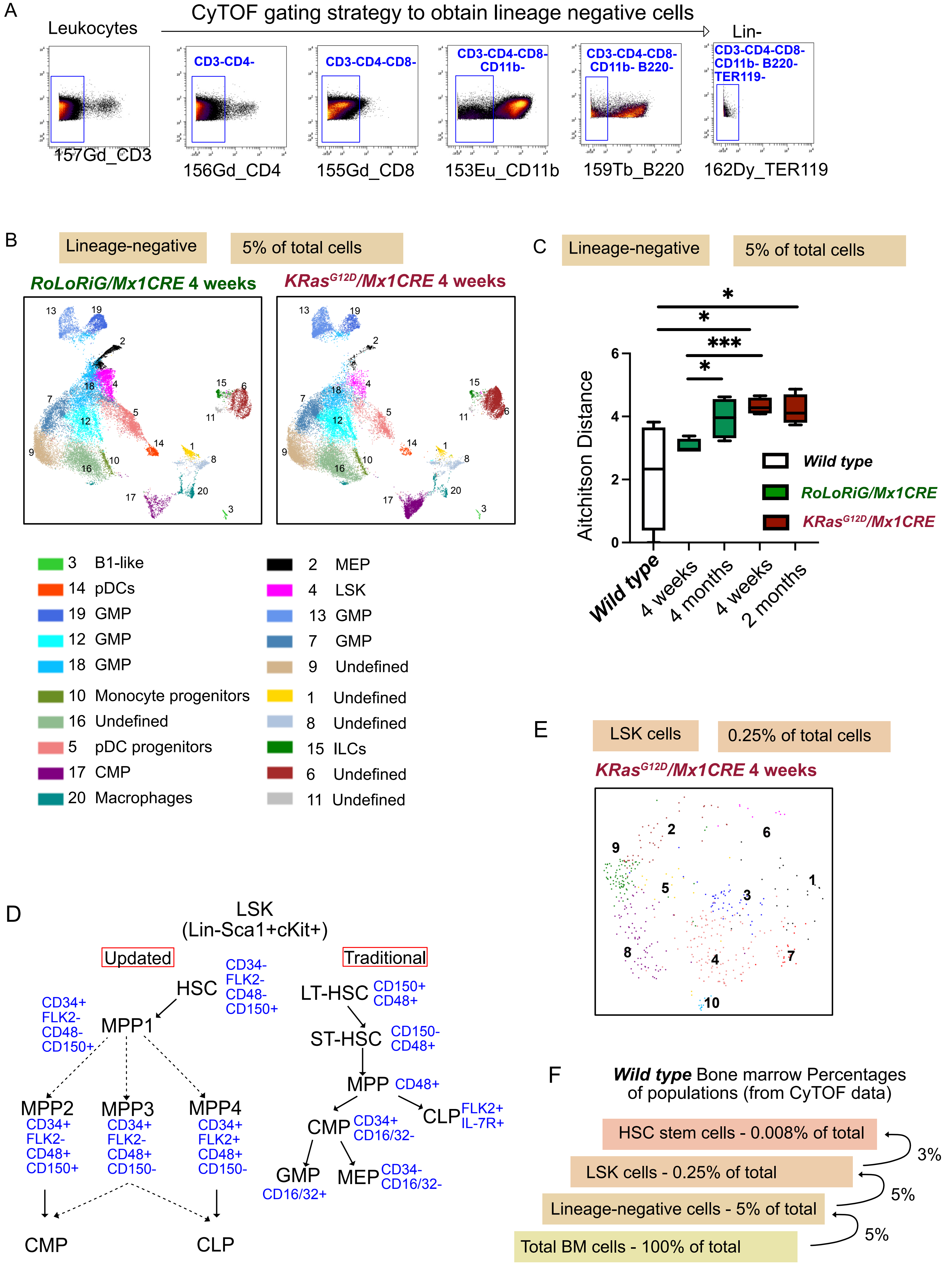

**Karra *et al.* , Supplemental Figure S8**

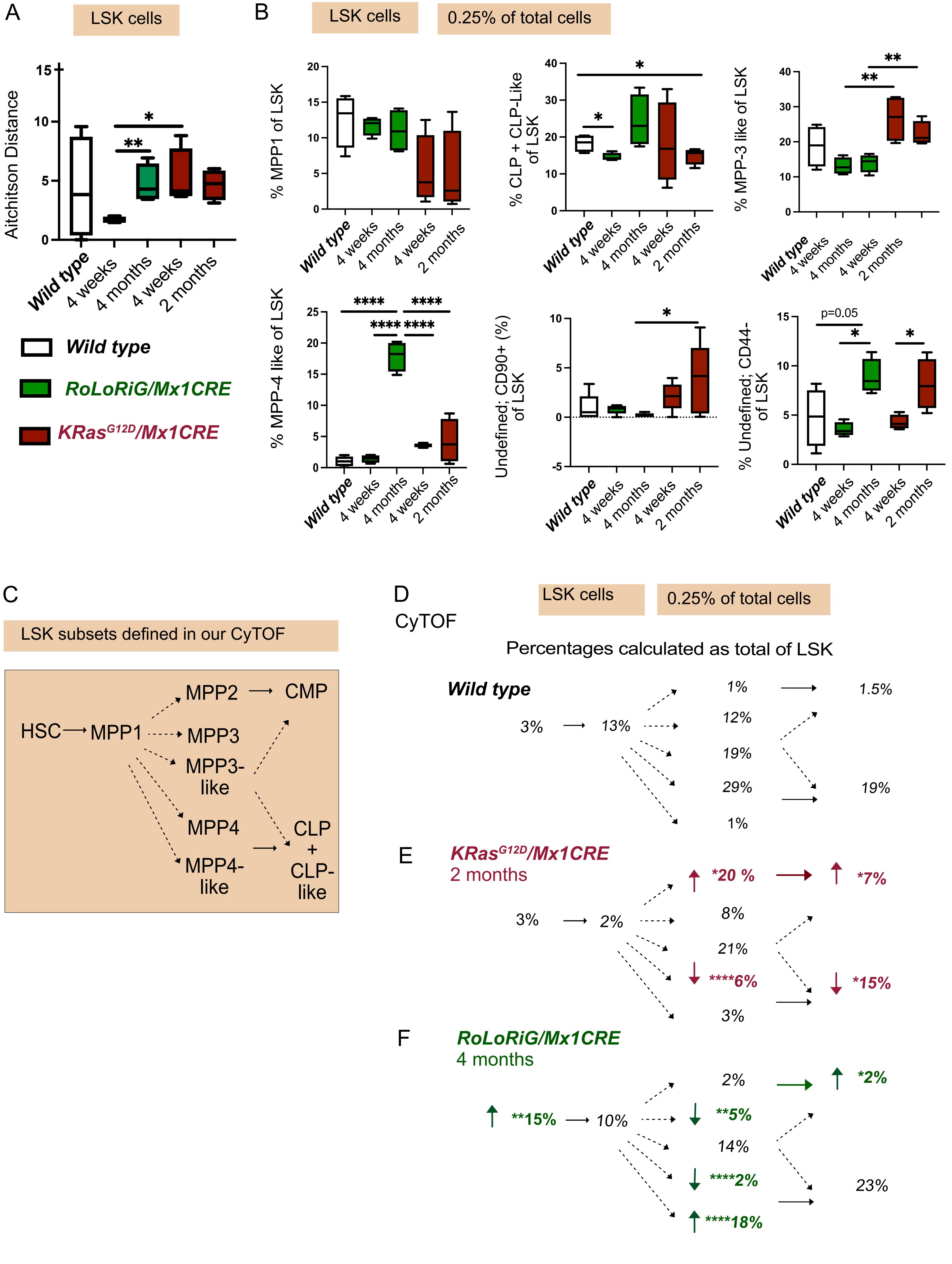

Karra *et al.* , Supplemental Figure S9

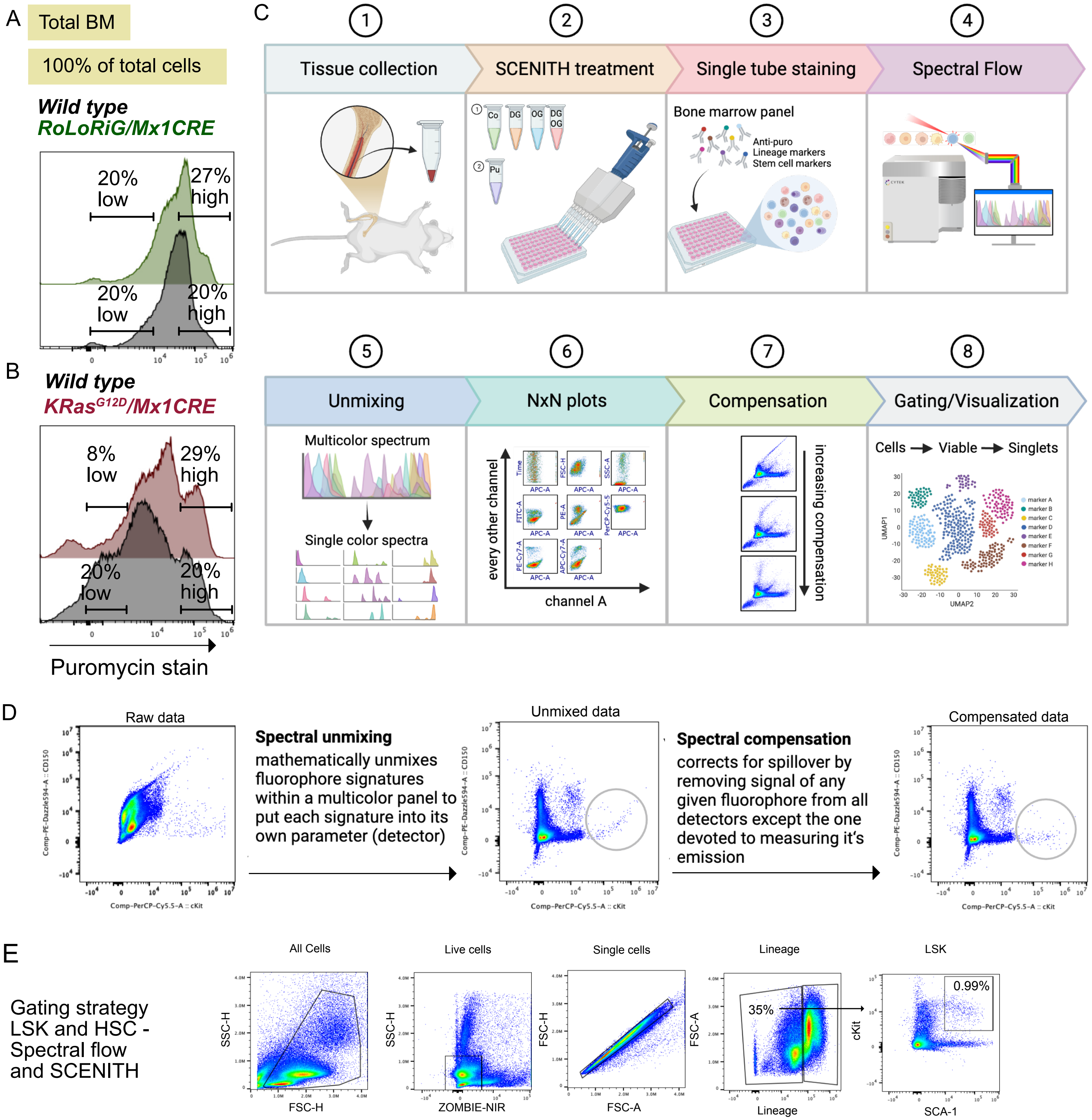

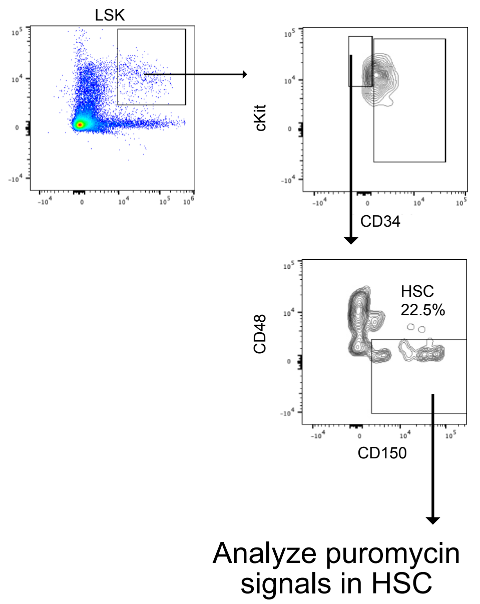

F

HSC numbers → Relative HSC frequency WT arbitrarily set to 20%

| *Wild type* | *KRasG12D/Mx1CRE* | *Wild type* | *KRasG12D/Mx1CRE* |
| --- | --- | --- | --- |
| 25.7% | 18.1% | 20.0% | 14.09% |
| | 6.52% | 20.0% | 5.08% |
| 27.3% | 10.5% | 20.0% | 7.70% |
| 16.3% | 6.06% | 20.0% | 7.44% |
| ***Wild type*** | ***RoLoRiG/Mx1CRE*** | ***Wild type*** | ***RoLoRiG/Mx1CRE*** |
| 22.5% | 16.8% | 20.0% | 14.93% |
| | 17.9% | 20.0% | 15.91% |
| 9.38% | 18.3% | 20.0% | 39.01% |
| | 13.8% | 20.0% | 29.42% |
| 19.4% | 27.8% | 20.0% | 28.70% |

**Karra *et al.* , Supplemental Figure S10**

Additional individual experiments (tSNE) - SCENITH on lineage-negative cells

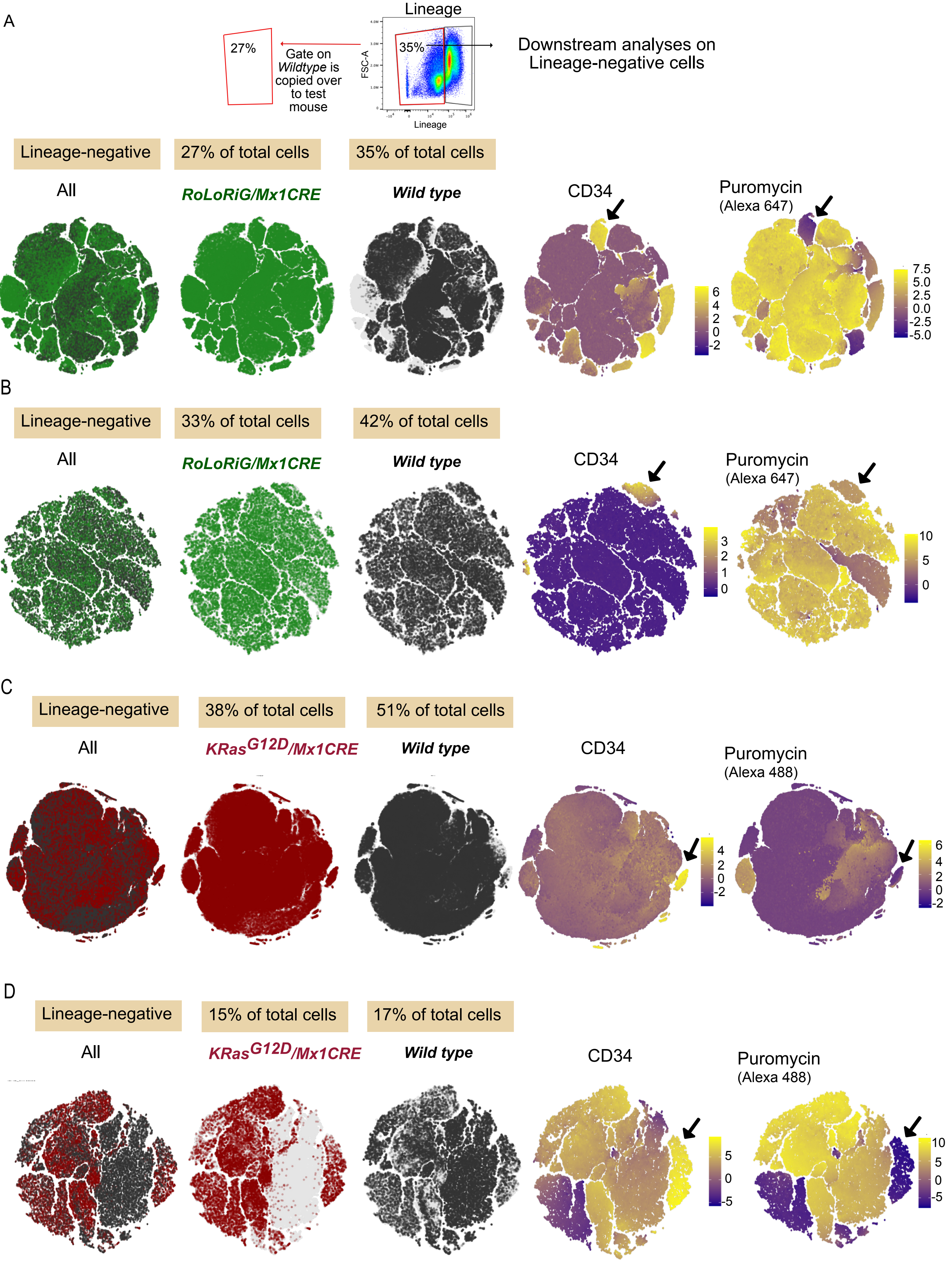

**Karra *et al.* , Supplemental Figure S11**

### Gating strategy MPP1, MPP2, MPP3, and MPP4

#### Spectral flow and SCENITH

Gating Strategy for MPPs

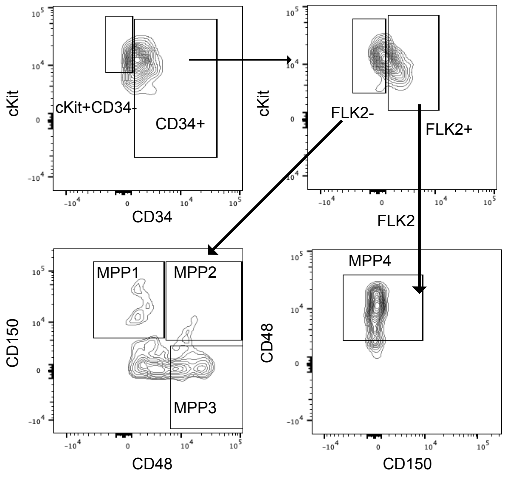

**Experimental set-up and gating strategy:**

Control *Wild type* mice and test mice were always stained in pairs in our experimental set-up.

Gates drawn for *Wild type* mice in the pair were copied over identically onto test mice.

MPP1 (CD34+FLK2-) → Relative MPP1 frequency WT arbitrarily set to 15

| *Wild type* | *$KRas^{G12D}$/Mx1CRE* | *Wild type* | *$KRas^{G12D}$/Mx1CRE* |
|---|---|---|---|
| 16.7% | 22.2% | 15.0% | 19.94% |
| | 22.6% | | 20.30% |
| 19.1% | 14.7% | 15.0% | 11.54% |
| 24% | 13.5% | 15.0% | 8.44% |

| *Wild type* | *RoLoRiG/Mx1CRE* | *Wild type* | *RoLoRiG/Mx1CRE* |
|---|---|---|---|
| 14.0% | 11.2% | 15.0% | 12.0% |
| | 13.9% | 15.0% | 14.89% |
| 10.6% | 15.3% | 15.0% | 21.65% |
| | 16.9% | 15.0% | 23.91% |
| 30.8% | 34.3% | 15.0% | 16.70% |

MPP2 (CD34+FLK2-) → Relative MPP2 frequency WT arbitrarily set to 20 and 8%

| *Wild type* | *$KRas^{G12D}$/Mx1CRE* | *Wild type* | *$KRas^{G12D}$/Mx1CRE* |
|---|---|---|---|
| 13.3% | 38.8% | 8.0% | 23.40% |
| | 48.4% | | 29.11% |
| 10.6% | 29.4% | 8.0% | 22.18% |
| 16.0% | 32.4% | 8.0% | 16.20% |

| *Wild type* | *RoLoRiG/Mx1CRE* | *Wild type* | *RoLoRiG/Mx1CRE* |
|---|---|---|---|
| 7.32% | 7.5% | 8.0% | 8.19% |
| | 16.0% | 8.0% | 17.48% |
| 6.62% | 16.6% | 8.0% | 20.06% |
| | 9.04% | 8.0% | 10.92% |
| 2.56% | 2.86% | 8.0% | 8.93% |

MPP3 (CD34+FLK2-) → Relative MPP3 frequency WT arbitrarily set to 40%

| *Wild type* | *$KRas^{G12D}$/Mx1CRE* | *Wild type* | *$KRas^{G12D}$/Mx1CRE* |
|---|---|---|---|
| 42.2% | 33.32% | 40.0% | 31.60% |
| | 29.0% | | 27.49% |
| 63.8% | 50.0% | 40.0% | 31.34% |
| 48.6% | 48.0% | 40.0% | 39.50% |

| *Wild type* | *RoLoRiG/Mx1CRE* | *Wild type* | *RoLoRiG/Mx1CRE* |
|---|---|---|---|
| 32.3% | 45.4% | 40.0% | 56.22% |
| | 38.5% | 40.0% | 47.67% |
| 45.7% | 47.8% | 40.0% | 41.83% |
| | 45.2% | 40.0% | 39.56% |
| 28.2% | 14.3% | 40.0% | 20.28% |

MPP4 (CD34+FLK2+) → Relative MPP4 frequency WT arbitrarily set to 80%

| *Wild type* | *$KRas^{G12D}$/Mx1CRE* | *Wild type* | *$KRas^{G12D}$/Mx1CRE* |
|---|---|---|---|
| 76.4% | 75.0% | 80.0% | 78.53% |
| | 74.0% | | 77.49% |
| 92.2% | 84.6% | 80.0% | 73.40% |
| 91.7% | 94.4% | 80.0% | 82.35% |

| *Wild type* | *RoLoRiG/Mx1CRE* | *Wild type* | *RoLoRiG/Mx1CRE* |
|---|---|---|---|
| 85.3% | 86.0% | 80.0% | 80.65% |
| | 79.8% | 80.0% | 74.84% |
| 90.7% | 75.3% | 80.0% | 66.41% |
| | 73.7% | 80.0% | 65.00% |
| 74.0% | 60.6% | 80.0% | 65.51% |

**Supplemental Table S1 Mass cytometry antibody panel**

| Channel | Metal | Antibody | Clone | Vendor | Catalogue number |
|---|---|---|---|---|---|
| 113 | In | *CD45.1* | | Biolegend | 110702 |
| 115 | In | *CD45.2* | | Biolegend | 109843 |
| 139 | La | Ly6G | 1A8 | Biolegend | 127602 |
| 140 | Ce | IgD | 11-26c.2a | BD Biosciences | 553438 |
| 141 | Pr | CD16/32 | | BD Biosciences | 553142 |
| 142 | Nd | CD49b | HMα2 | Biolegend | 103513 |
| 143 | Nd | CD11c | HL3 | BD Biosciences | 553799 |
| 144 | Nd | CD43 | S7 | BD Biosciences | 553268 |
| 145 | Nd | CD27 | LG.3A10 | Biolegend | 124202 |
| 146 | Nd | CD138 | 281-2 | Biolegend | 142502 |
| 147 | Sm | CD34 | | BD Biosciences | 553731 |
| 148 | Nd | CD41 | MwReg30 | BD Biosciences | 553847 |
| 149 | Sm | SiglecF | E50-2440 | BD Biosciences | 552125 |
| 150 | Nd | PDCA1 | 120g8 | Novus Biologicals | DDX0390-HD05 |
| 151 | Eu | Ly6C | HK1.4 | Biolegend | 128001 |
| 152 | Sm | CD48 | | BD Biosciences | 555758 |
| 153 | Eu | CD11b | M1/70 | Biolegend | 101202 |
| 154 | Sm | cKit | 2B8 | Biolegend | 105802 |
| 155 | Gd | CD8 | 53-6.7 | Biolegend | 100702 |
| 156 | Gd | CD4 | RM4-5 | Biolegend | 100506 |
| 157 | Gd | CD3 | 17A2 | BD Biosciences | 555273 |
| 158 | Gd | TCRb | **H57-597** | Biolegend | 109202 |
| 159 | Tb | B220 | 188 | Biolegend | 103202 |
| 161 | Dy | CD127 | SB/199 | BIolegend | 121102 |

| | | | | | |
|---|---|---|---|---|---|
| 162 | Dy | *Ter119* | TER119 | Fluidigm | 3162003B |
| 163 | Dy | CD105 | MJ7/15 | Biolegend | 120402 |
| 164 | Dy | Sca1 | D7 | Biolegend | 108102 |
| 165 | Ho | CD150 | 9D1 | Novus biologicals | NB100-63446 |
| 166 | Er | FcER1a | MAR-1 | Biolegend | 134321 |
| 167 | Er | CD135 | A2F10 | Peprotech | 17412-20 |
| 168 | Er | CD25 | 3C7 | Biolegend | 101902 |
| 169 | Tm | F4/80 | BM8 | Biolegend | 123143 |
| 170 | Er | CD115 | AFS98 | Biolegend | 135502 |
| 171 | Yb | CD71 | R17217 | Biolegend | 113802 |
| 172 | Yb | CD23 | B3B4 | Biolegend | 101602 |
| 173 | Yb | CD19 | 6D5 | Biolegend | 115547 |
| 174 | Yb | IgM | RMM-1 | Biolegend | 406502 |
| 175 | Lu | CD44 | IM7 | BD Biosciences | 553131 |
| 176 | Yb | CD90 | G7 | Biolegend | 105202 |
| 209 | Bi | MHC II | M5/114.15.2 | Biolegend | 107602 |

| **Cluster number** | **Markers expressed (analysis in Total BM Figure 1)** |
|---|---|
| 3 | $MHCII^{low}$ $CD44^{+}$ $IgM^{low}$ $CD150^{+}$ $TCRb^{low}$ $CD4^{+}$ $CD8^{+}$ $ckit^{low}$ $CD11b^{+}$ $CD48^{med}$ $Ly6C^{+}$ $PDCA1^{low}$ $CD41^{low}$ $CD34^{low}$ $CD138^{low}$ $CD27^{low}$ $CD43^{+}$ $CD11c^{low}$ $CD49b^{+}$ $CD16/32^{+}$ $IgD^{+}$ $Ly6G^{+}$ |
| 15 | $CD44^{med}$ $CD71^{med}$ $F480^{low}$ $CD25^{low}$ $SCA1^{low}$ $CD105^{low}$ $CD3^{low}$ $CD4^{low}$ $CD8^{low}$ $cKit^{+}$ $CD11b^{low}$ $CD48^{+}$ $Ly6C^{+}$ $PDCA1^{low}$ $SiglecF^{low}$ $CD41^{low}$ $CD34^{+}$ $CD27^{low}$ $CD43^{med}$ $CD16/32^{+}$ $Ly6G^{med}$ |
| 16 | $CD3^{low}$ $CD8^{low}$ $cKit^{low}$ $CD11B^{+}$ $CD48^{med}$ $Ly6C^{+}$ $SiglecF^{low}$ $CD34^{low}$ $CD43^{+}$ $CD16/32^{+}$ $IgD^{low}$ $Ly6G^{+}$ |
| 22 | $MHCII^{low}$ $CD90^{+}$ $CD44^{+}$ $IgM^{low}$ $CD71^{low}$ $F480^{low}$ $CD25^{low}$ $CD150^{low}$ $SCA1^{med}$ $B220^{low}$ $TCRb^{+}$ $CD3^{+}$ $CD4^{med}$ $CD8^{med}$ $cKit^{low}$ $CD11b^{+}$ $CD48^{+}$ $Ly6C^{+}$ $PDCA1^{low}$ $SiglecF^{low}$ $CD41^{low}$ $CD34^{low}$ $CD27^{low}$ $CD43^{+}$ $CD11C^{low}$ $CD49b^{low}$ $CD16/32^{+}$ $IgD^{low}$ $Ly6G^{+}$ |
| 23 | $MHCII^{low}$ $CD90^{low}$ $CD44^{+}$ $CD71^{low}$ $F4/80^{med}$ $CD25^{low}$ $SCA1^{low}$ $B220^{low}$ $TCRb^{low}$ $CD3^{low}$ $CD4^{low}$ $CD8^{med}$ $cKit^{low}$ $CD11b^{+}$ $CD48^{+}$ $Ly6C^{+}$ $PDCA1^{med}$ $SiglecF^{low}$ $CD41^{low}$ $CD34^{med}$ $CD27^{low}$ $CD43^{+}$ $CD16/32^{+}$ $IgD^{low}$ $Ly6G^{+}$ |
| 25 | $MHCII^{+}$ $CD44^{+}$ $IgM^{med}$ $CD19^{med}$ $CD71^{+}$ $F4/80^{low}$ $CD25^{low}$ $SCA1^{low}$ $B220^{+}$ $CD3^{low}$ $CD4^{low}$ $CD8^{med}$ $cKit^{low}$ $CD11b^{+}$ $CD48^{+}$ $Ly6C^{+}$ $PDCA1^{low}$ $SiglecF^{low}$ $CD34^{low}$ $CD43^{+}$ $CD16/32^{+}$ $IgD^{low}$ $Ly6G^{+}$ |

**Karra *et al.* , Supplemental Table 2**

| Cluster number | Population name | Makers expressed (Lineage negative cells in Figure 3 ) |
|---|---|---|
| 1 | Undefined | MHCII$^{+}$ CD44$^{low}$ IgM$^{low}$ CD19$^{low}$ CD71$^{+}$ CD48$^{+}$ Ly6C$^{+}$ CD43$^{low}$ CD16/32$^{+}$ Ly6G$^{+}$ |
| 2 | MEP | CD44$^{+}$ CD23$^{low}$ CD71$^{+}$ F4/80$^{med}$ CD25$^{low}$ CD150$^{med}$ SCA1$^{low}$ CD105$^{+}$ cKit$^{+}$ CD48$^{+}$ Ly6C$^{+}$ PDCA1$^{med}$ SiglecF$^{med}$ CD41$^{med}$ CD34$^{+}$ CD27$^{low}$ CD43$^{+}$ CD16/32$^{+}$ Ly6G$^{+}$ |
| 3 | B1-like | MHCII$^{+}$ CD44$^{+}$ IgM$^{+}$ CD19$^{+}$ CD23$^{low}$ CD71$^{med}$ F4/80$^{low}$ SCA1$^{+}$ cKit$^{low}$ CD48$^{+}$ Ly6C$^{+}$ PDCA1$^{low}$ CD34$^{low}$ CD43$^{+}$ CD16/32$^{+}$ IgD$^{low}$ Ly6G$^{+}$ |
| 4 | LSK | MHCII$^{low}$ CD44$^{+}$ CD71$^{low}$ F4/80$^{low}$ CD25$^{low}$ CD135$^{med}$ SCA1$^{+}$ CD105$^{med}$ cKit$^{+}$ CD48$^{+}$ Ly6C$^{+}$ PDCA1$^{med}$ CD41$^{med}$ CD34$^{+}$ CD138$^{low}$ CD27$^{+}$ CD43$^{+}$ CD49b$^{low}$ CD16/32$^{+}$ Ly6G$^{+}$ |
| 5 | pDC progenitors | MHCII$^{low}$ CD44$^{+}$ CD71$^{+}$ CD25$^{low}$ CD135$^{low}$ FcERI$^{low}$ SCA1$^{low}$ CD105$^{low}$ cKit$^{low}$ CD48$^{+}$ Ly6C$^{+}$ PDCA1$^{+}$ CD41$^{low}$ CD34$^{+}$ CD138$^{low}$ CD27$^{med}$ CD43$^{+}$ CD11c$^{+}$ CD16/32$^{+}$ Ly6G$^{+}$ |
| 6 | Undefined | CD90$^{+}$ CD44$^{+}$ CD48$^{+}$ Ly6C$^{med}$ CD27$^{med}$ CD43$^{med}$ CD11c$^{med}$ CD49b$^{+}$ CD16/32$^{+}$ Ly6G$^{med}$ |
| 7 | GMP | CD90$^{low}$ CD44$^{+}$ CD23$^{low}$ CD71$^{+}$ CD115$^{med}$ F4/80$^{med}$ CD25$^{med}$ CD135$^{med}$ SCA1$^{low}$ CD105$^{med}$ cKit$^{+}$ CD48$^{+}$ Ly6C$^{+}$ PDCA1$^{+}$ SiglecF$^{med}$ CD41$^{low}$ CD34$^{+}$ CD27$^{med}$ CD43$^{+}$ CD16/32$^{+}$ Ly6G$^{+}$ |
| 8 | Undefined | MHCII$^{med}$ CD44$^{med}$ CD48$^{low}$ Ly6C$^{low}$ CD43$^{low}$ CD16/32$^{med}$ IgD$^{med}$ Ly6G$^{low}$ |
| 9 | Undefined | CD90$^{low}$ CD44$^{+}$ CD71$^{+}$ CD115$^{med}$ F4/80$^{+}$ CD25$^{low}$ CD135$^{low}$ SCA1$^{low}$ CD105$^{low}$ cKit$^{low}$ CD48$^{+}$ Ly6C$^{+}$ PDCA1$^{med}$ SiglecF$^{low}$ CD41$^{low}$ CD34$^{+}$ CD43$^{med}$ CD16/32$^{+}$ Ly6G$^{+}$ |
| 10 | Monocyte progenitors | CD71$^{low}$ CD48$^{+}$ Ly6C$^{+}$ PDCA1$^{low}$ CD34$^{+}$ CD43$^{med}$ CD16/32$^{+}$ Ly6G$^{+}$ |
| 11 | Undefined | CD90$^{+}$ CD44$^{low}$ TCRb$^{+}$ CD48$^{+}$ Ly6C$^{+}$ CD27$^{med}$ CD43$^{+}$ CD49b$^{low}$ CD16/32$^{low}$ Ly6G$^{med}$ |
| 12 | GMP | CD44$^{med}$ CD71$^{+}$ cKit$^{+}$ CD48$^{+}$ Ly6C$^{+}$ PDCA1$^{med}$ SiglecF$^{low}$ CD34$^{+}$ CD27$^{low}$ CD43$^{med}$ CD16/32$^{+}$ Ly6G$^{+}$ |
| 13 | GMP | CD44$^{+}$ CD71$^{+}$ F4/80$^{+}$ CD25$^{low}$ cKit$^{med}$ CD48$^{+}$ Ly6C$^{+}$ PDCA1$^{low}$ SiglecF$^{+}$ CD41$^{med}$ CD34$^{+}$ CD43$^{+}$ CD16/32$^{+}$ IgD$^{low}$ Ly6G$^{+}$ |
| 14 | pDCs | MHCII$^{+}$ CD44$^{+}$ CD25$^{low}$ SCA1$^{+}$ CD105$^{low}$ cKit$^{low}$ CD48$^{+}$ Ly6C$^{+}$ PDCA1$^{+}$ CD34$^{low}$ CD138$^{low}$ CD43$^{+}$ CD16/32$^{+}$ Ly6G$^{+}$ |
| 15 | ILCs | MHCII$^{med}$ CD90$^{+}$ CD44$^{+}$ F4/80$^{med}$ CD25$^{+}$ SCA1$^{+}$ CD105$^{low}$ CD127$^{low}$ cKit$^{low}$ CD48$^{+}$ Ly6C$^{+}$ CD27$^{med}$ CD43$^{med}$ CD49b$^{low}$ CD16/32$^{+}$ Ly6G$^{+}$ |
| 16 | Undefined | CD44$^{+}$ CD71$^{med}$ F4/80$^{low}$ CD48$^{+}$ Ly6C$^{+}$ PDCA1$^{med}$ CD34$^{+}$ CD43$^{med}$ CD16/32$^{+}$ Ly6G$^{+}$ |
| 17 | CMP | MHCII$^{+}$ CD44$^{+}$ CD71$^{+}$ F4/80$^{low}$ CD25$^{low}$ CD135$^{low}$ SCA1$^{low}$ cKit$^{+}$ CD48$^{+}$ Ly6C$^{+}$ PDCA1$^{med}$ SiglecF$^{low}$ CD41$^{low}$ CD34$^{+}$ CD43$^{+}$ CD11c$^{+}$ CD16/32$^{+}$ Ly6G$^{+}$ |
| 18 | GMP | CD44$^{+}$ CD71$^{+}$ F4/80$^{low}$ CD25$^{low}$ CD105$^{low}$ cKit$^{+}$ CD48$^{+}$ Ly6C$^{+}$ PDCA1$^{med}$ SiglecF$^{med}$ CD41$^{med}$ CD34$^{+}$ CD27$^{+}$ CD43$^{+}$ CD16/32$^{+}$ Ly6G$^{+}$ |
| 19 | GMP (Basophils/Mast cell progenitors) PMID: 24598075 | CD90$^{low}$ CD44$^{med}$ CD71$^{med}$ F4/80$^{low}$ CD25$^{low}$ CD135$^{low}$ FcERI$^{+}$ SCA1$^{low}$ cKit$^{low}$ CD48$^{+}$ Ly6C$^{+}$ PDCA1$^{low}$ SiglecF$^{med}$ CD41$^{+}$ CD34$^{+}$ CD43$^{+}$ CD11c$^{low}$ CD49b$^{+}$ CD16/32$^{+}$ Ly6G$^{+}$ |
| 20 | Macrophages | MHCII$^{+}$ CD44$^{low}$ CD71$^{+}$ F4/80$^{+}$ SCA1$^{low}$ CD48$^{+}$ Ly6C$^{+}$ PDCA1$^{low}$ CD41$^{low}$ CD34$^{low}$ CD43$^{low}$ CD11c$^{low}$ CD16/32$^{+}$ IgD$^{low}$ Ly6G$^{+}$ |

| Cluster number | Population name | Markers expressed (LSK analysis in Figure 4 ) |
|---|---|---|
| 1 | MPP1 | MHCII$^{med}$ CD44$^{med}$ F4/80$^{low}$ SCA1$^{+}$CD105$^{low}$ cKit$^{+}$CD48$^{+}$ Ly6C$^{+}$PDCA1$^{low}$ CD41$^{low}$ CD34$^{+}$ CD27$^{+}$CD43$^{+}$CD49b$^{med}$ CD16/32$^{+}$Ly6G$^{+}$ |
| 2 | MPP4 | MHCII$^{low}$ CD90$^{low}$ CD44$^{+}$CD71$^{+}$F4/80$^{+}$ CD25$^{low}$ CD135$^{+}$ SCA1$^{+}$CD105$^{+}$ cKit$^{+}$ CD48$^{+}$ Ly6C$^{+}$PDCA1$^{med}$ SiglecF$^{low}$ CD41$^{med}$ CD34$^{+}$CD27$^{+}$CD43$^{+}$ CD49b$^{med}$ CD16/32$^{+}$ Ly6G$^{+}$ |
| 3 | MPP3 | MHCII$^{low}$ CD44$^{+}$CD71$^{low}$ F4/80$^{low}$ CD135$^{low}$ SCA1$^{+}$CD105$^{low}$ cKit$^{+}$CD48$^{+}$Ly6C$^{+}$ PDCA1$^{med}$ SiglecF$^{low}$ CD41$^{med}$ CD34$^{+}$CD27$^{+}$CD43$^{+}$CD49b$^{med}$ CD16/32$^{+}$ Ly6G$^{+}$ |
| 4 | MPP3-like | MHCII$^{low}$ CD44$^{+}$CD71$^{med}$ CD135$^{med}$ SCA1$^{+}$ CD105$^{low}$ cKit$^{+}$CD48$^{+}$Ly6C$^{+}$ PDCA1$^{med}$ CD41$^{low}$ CD34$^{+}$CD27$^{+}$CD43$^{+}$ CD16/32$^{+}$ Ly6G$^{+}$ |
| 5 | MPP4-like | MHCII$^{+}$CD44$^{+}$CD71$^{med}$ F4/80$^{low}$ CD25$^{low}$ CD135$^{med}$ CD150$^{low}$ SCA1$^{+}$ CD105$^{+}$cKit$^{+}$ CD48$^{+}$Ly6C$^{+}$ PDCA1$^{med}$ SiglecF$^{med}$ CD41$^{med}$ CD34$^{+}$ CD27$^{+}$CD43$^{+}$ CD49b$^{low}$ CD16/32$^{+}$ Ly6G$^{+}$ |
| 6 | HSC | MHCII$^{med}$ CD90$^{low}$ CD44$^{+}$ CD71$^{low}$ F4/80$^{+}$CD25$^{low}$ CD150$^{+}$SCA1$^{+}$ CD105$^{+}$ cKit$^{+}$ CD48$^{+}$Ly6C$^{+}$ PDCA1$^{med}$ CD41$^{med}$ CD34$^{+}$ CD27$^{med}$ CD43$^{+}$CD49b$^{low}$ CD16/32$^{+}$Ly6G$^{+}$ |
| 7 | Undefined | MHCII$^{med}$ CD71$^{low}$ CD135$^{low}$ SCA1$^{med}$ CD105$^{low}$ cKit$^{+}$CD48$^{+}$ Ly6C$^{+}$PDCA1$^{med}$ CD41$^{low}$ CD34$^{+}$CD27$^{+}$CD43$^{+}$CD16/32$^{+}$Ly6G$^{+}$ |
| 8 | CLP-like | MHCII$^{low}$ CD90$^{low}$ CD44$^{+}$CD71$^{+}$F4/80$^{low}$ CD25$^{low}$ CD135$^{+}$SCA1$^{+}$CD105$^{med}$ CD127$^{low}$ cKit$^{+}$CD48$^{+}$ Ly6C$^{+}$PDCA1$^{med}$ CD41$^{low}$ CD34$^{+}$ CD27$^{+}$CD43$^{+}$ CD16/32$^{+}$ Ly6G$^{+}$ |
| 9 | MPP2 | MHCII$^{low}$ CD44$^{+}$IgM$^{med}$ CD23$^{low}$ CD71$^{+}$F4/80$^{med}$ CD25$^{+}$CD135$^{+}$FcERI$^{low}$ CD150$^{med}$ SCA1$^{+}$ CD105$^{+}$TCRb$^{med}$ cKit$^{+}$CD48$^{+}$ Ly6C$^{+}$ PDCA1$^{med}$ SiglecF$^{med}$ CD41$^{med}$ CD34$^{+}$ CD138$^{low}$ CD27$^{+}$ CD43$^{+}$ CD11c$^{low}$ CD16/32$^{+}$ IgD$^{low}$ Ly6G$^{+}$ |
| 10 | Undefined | MHCII$^{low}$ CD90$^{+}$CD44$^{+}$IgM$^{low}$ CD71$^{med}$ CD25$^{low}$ SCA1$^{med}$ CD105$^{+}$CD127$^{low}$ TCRb$^{low}$ cKit$^{+}$CD48$^{+}$ Ly6C$^{+}$CD41$^{low}$CD34$^{+}$CD27$^{+}$CD43$^{+}$CD16/32$^{med}$ Ly6G$^{+}$ |

**Karra *et al.* , Supplemental Table 4**

#### KEY RESOURCES TABLE

The table highlights the reagents, genetically modified organisms and strains, cell lines, software, instrumentation, and source data **essential** to reproduce results presented in the manuscript. Depending on the nature of the study, this may include standard laboratory materials (i.e., food chow for metabolism studies, support material for catalysis studies), but the table is **not** meant to be a comprehensive list of all materials and resources used (e.g., essential chemicals such as standard solvents, SDS, sucrose, or standard culture media do not need to be listed in the table). **Items in the table must also be reported in the method details section within the context of their use.** To maximize readability, the number of **oligonucleotides and RNA sequences** that may be listed in the table is restricted to no more than 10 each. If there are more than 10 oligonucleotides or RNA sequences to report, please provide this information as a supplementary document and reference the file (e.g., See Table S1 for XX) in the key resources table.

***Please note that ALL references cited in the key resources table must be included in the references list.*** Please report the information as follows:

- **REAGENT or RESOURCE:** Provide full descriptive name of the item so that it can be identified and linked with its description in the manuscript (e.g., provide version number for software, host source for antibody, strain name). In the experimental models section (applicable only to experimental life science studies), please include all models used in the paper and describe each line/strain as: model organism: name used for strain/line in paper: genotype. (i.e., Mouse: OXTR$^{fl/fl}$: B6.129(SJL)-Oxtr$^{tm1.1Wsy/J}$). In the biological samples section (applicable only to experimental life science studies), please list all samples obtained from commercial sources or biological repositories. Please note that software mentioned in the methods details or data and code availability section needs to also be included in the table. See the sample tables at the end of this document for examples of how to report reagents.

- **SOURCE:** Report the company, manufacturer, or individual that provided the item or where the item can be obtained (e.g., stock center or repository). For materials distributed by Addgene, please cite the article describing the plasmid and include "Addgene" as part of the identifier. If an item is from another lab, please include the name of the principal investigator and a citation if it has been previously published. If the material is being reported for the first time in the current paper, please indicate as "this paper." For software, please provide the company name if it is commercially available or cite the paper in which it has been initially described.

- **IDENTIFIER:** Include catalog numbers (entered in the column as "Cat#" followed by the number, e.g., Cat#3879S). Where available, please include unique entities such as RRIDs, Model Organism Database numbers, accession numbers, and PDB, CAS, or CCDC IDs. For antibodies, if applicable and available, please also include the lot number or clone identity. For software or data resources, please include the URL where the resource can be downloaded. Please ensure accuracy of the identifiers, as they are essential for generation of hyperlinks to external sources when available. Please see the Elsevier list of data repositories with automated bidirectional linking for details. When listing more than one identifier for the same item, use semicolons to separate them (e.g., Cat#3879S; RRID: AB_2255011). If an identifier is not available, please enter "N/A" in the column.
    - ***A NOTE ABOUT RRIDs:*** We highly recommend using RRIDs as the identifier (in particular for antibodies and organisms but also for software tools and databases). For more details on how to obtain or generate an RRID for existing or newly generated resources, please visit the RII or search for RRIDs.

Please use the empty table that follows to organize the information in the sections defined by the subheading, skipping sections not relevant to your study. Please do not add subheadings. To add a row, place the cursor at the end of the row above where you would like to add the row, just outside the right border of the table. Then press the ENTER key to add the row. Please delete empty rows. Each entry must be on a separate row; do not list multiple items in a single table cell. Please see the sample tables at the end of this document for relevant examples in the life and physical sciences of how reagents and instrumentation should be cited.

*TABLE FOR AUTHOR TO COMPLETE*

*Please upload the completed table as a separate document.* ***Please do not add subheadings to the key resources table.*** *If you wish to make an entry that does not fall into one of the subheadings below, please contact your handling editor.* ***Any subheadings not relevant to your study can be skipped.*** *(**NOTE:** For authors publishing in Cell Genomics, Cell Reports Medicine, Current Biology, and Med, please note that references within the KRT should be in numbered style rather than Harvard.)*

**KEY RESOURCES TABLE**

| REAGENT or RESOURCE | SOURCE | IDENTIFIER |
|---|---|---|
| CyTOF Antibodies (mouse) | | |
| Anti-mouse CD45.1 | Biolegend | Cat# 110702; **RRID:AB_313491** |
| Anti-mouse CD45.2 | Biolegend | Cat# 109843; **RRID:AB_2563751** |
| Anti-mouse Ly6G | Biolegend | Cat# 127602 Clone 1A8; **RRID:AB_1089180** |
| Anti-mouse IgD | BD Biosciences | Cat# 553438 Clone11-26c.2a; |
| Anti-mouse CD16/32 | BD Biosciences | Cat# 553142; **RRID:AB_394657** |
| Anti-mouse CD49b | Biolegend | Cat# 103513 Clone Hmα2; **RRID:AB_2563754** |
| Anti-mouse CD11c | BD Biosciences | Cat# 553799 Clone HL3; **RRID:AB_395058** |
| Anti-mouse CD43 | BD Biosciences | Cat# 553268 Clone S7; **RRID:AB_394745** |
| Anti-mouse CD27 | Biolegend | Cat# 124202 Clone LG.3A10; **RRID:AB_1236456** |
| Anti-mouse CD138 | Biolegend | Cat# 142502 Clone 281-2; **RRID:AB_10965646** |
| Anti-mouse CD34 | BD Biosciences | Cat# 553731; **RRID:AB_395015** |
| Anti-mouse CD41 | BD Biosciences | Cat# 553847 Clone MwReg30; **RRID:AB_395084** |

| | | |
|---|---|---|
| Anti-mouse Siglec-F | BD Biosciences | Cat# 552125 Clone E50-2440; **RRID:AB_394340** |
| Anti-mouse PDCA-1 | Novus Biologicals | Cat# DDX0390-HD05Clone 120g8 |
| Anti-mouse Ly6C | Biolegend | Cat# 128001 Clone HK1.4; **RRID:AB_1134213** |
| Anti-mouse CD48 | BD Biosciences | Cat# 555758; **RRID:AB_396099** |
| Anti-mouse CD11b | Biolegend | Cat# 101202 Clone M1/70; **RRID:AB_312785** |
| Anti-mouse cKit | Biolegend | Cat# 105802 Clone 2B8; **RRID:AB_313211** |
| Anti-mouse CD8 | Biolegend | Cat# 100702 Clone 53-6.7; **RRID:AB_312741** |
| Anti-mouse CD4 | Biolegend | Cat# 100506 Clone RM4-5; **RRID:AB_312709** |
| Anti-mouse CD3 | BD Biosciences | Cat#555273 Clone 17A2; **RRID:AB_395697** |
| Anti-mouse TCRb | Biolegend | Cat# 109202 Clone H57-597; **RRID:AB_313425** |
| Anti-mouse B220 | Biolegend | Cat# 103202 Clone 188; **RRID:AB_312987** |
| Anti-mouse CD127 | BIolegend | Cat#121102 Clone SB/199; **RRID:AB_493500** |
| Anti-mouse TER119 | Fluidigm | Cat# 3162003B Clone TER119; |
| Anti-mouse CD105 | Biolegend | Cat# 120402 Clone MJ7/15; **RRID:AB_961070** |
| Anti-mouse SCA1 | Biolegend | Cat# 108102 Clone D7; **RRID:AB_313339** |

| | | |
|---|---|---|
| Anti-mouse CD150 | Novus biologicals | Cat# NB100-63446 Clone 9D1; **RRID:AB_959730** |
| Anti-mouse FCERIa | Biolegend | Cat# 134321 Clone MAR-1; **RRID:AB_2563768** |
| Anti-mouse CD135 | Peprotech | Cat# 17412-20 Clone A2F10 |
| Anti-mouse CD25 | Biolegend | Cat# 101902 Clone 3C7; **RRID:AB_312845** |
| Anti-mouse F4/80 | Biolegend | Cat# 123143 Clone BM8; **RRID:AB_2563767** |
| Anti-mouse CD115 | Biolegend | Cat# 135502 Clone AFS98; **RRID:AB_1937293** |
| Anti-mouse CD71 | Biolegend | Cat# 113802 Clone R17217; **RRID:AB_313563** |
| Anti-mouse CD23 | Biolegend | Cat# 101602 Clone B3B4; **RRID:AB_312827** |
| Anti-mouse CD19 | Biolegend | Cat# 115547 Clone 6D5; **RRID:AB_2562806** |
| Anti-mouse IgM | Biolegend | Cat# 406502 Clone RMM-1; **RRID:AB_315052** |
| Anti-mouse CD44 | BD Biosciences | Cat# 553131 Clone IM7; **RRID:AB_394646** |
| Anti-mouse CD90 | Biolegend | Cat# 105202 Clone G7; **RRID:AB_313169** |
| Anti-mouse MHCII | Biolegend | Cat# 107602 Clone M5/114.15; **RRID:AB_313317** |
| Flow Cytometry antibodies (mouse) | | |
| Anti-mouse PE-Cy™7 Rat Anti-Mouse CD3 Molecular Complex | BD Biosciences | Cat#5 60591; RRID:AB_1727462 |

| | | |
|---|---|---|
| Anti-mouse PE-Cy™7 Rat Anti-Mouse TER-119/Erythroid Cells | BD Biosciences | Cat# 557853; RRID:AB_396898 |
| Anti-mouse CD11b Monoclonal Antibody (M1/70), PE-Cyanine7 | eBioscience | Cat# 25-0112-82; RRID:AB_469588 |
| Anti-mouse PE/Cyanine7 anti-mouse CD5 Antibody | Biolegend | Cat# 100621; RRID:AB_2562772 |
| Anti-mouse CD45R (B220) Monoclonal Antibody (RA3-6B2), PE-Cyanine5.5 | eBioscience | Cat# 35-0452-82; RRID:AB_469721 |
| Anti-mouse CD4 Monoclonal Antibody (RM4-5), PE-Cyanine7 | eBioscience | Cat# 25-0042-82; RRID:AB_469578) |
| Anti-mouse CD8a Monoclonal Antibody (53-6.7), PE-Cyanine7 | eBioscience | Cat# 25-0081-82; RRID:AB_469584 |
| Anti-mouse CD117 Monoclonal Antibody (2B8) PerCP-Cy5.5 | BioLegend | Cat# 105824 RRID: AB_2131597 |
| Anti-mouse PE-Cy™7 Rat Anti-Mouse CD3 Molecular Complex | BD Biosciences | Cat#5 60591; RRID:AB_1727462 |
| Anti-mouse PE-Cy™7 Rat Anti-Mouse TER-119/Erythroid Cells | BD Biosciences | Cat# 557853; RRID:AB_396898 |
| Anti-mouse CD11b Monoclonal Antibody (M1/70), PE-Cyanine7 | eBioscience | Cat# 25-0112-82; RRID:AB_469588 |
| Anti-mouse PE/Cyanine7 anti-mouse CD5 Antibody | Biolegend | Cat# 100621; RRID:AB_2562772 |
| Anti-mouse CD45R (B220) Monoclonal Antibody (RA3-6B2), PE-Cyanine5.5 | eBioscience | Cat# 35-0452-82; RRID:AB_469721 |
| Anti-mouse CD4 Monoclonal Antibody (RM4-5), PE-Cyanine7 | eBioscience | Cat# 25-0042-82; RRID:AB_469578) |
| Anti-mouse CD8a Monoclonal Antibody (53-6.7), PE-Cyanine7 | eBioscience | Cat# 25-0081-82; RRID:AB_469584 |
| Anti-mouse CD117 Monoclonal Antibody (2B8) PerCP-Cy5.5 | BioLegend | Cat# 105824 RRID: AB_2131597 |
| Anti-mouse Ly-6A/Ly-6E Monoclonal Antibody Brilliant Violet 480 (E13-161.7) | BD Biosciences | Cat# 746546 RRID: AB_2743837 |
| Anti-mouse CD48 Monoclonal Antibody Brilliant Violet 605 (HM48-1) | BioLegend | Cat# 103441 RRID: AB_2650825 |
| Anti-mouse CD127 Monoclonal Antibody Brilliant Violet 421 (A7R34) | BioLegend | Cat# 135024 RRID: AB_11218800 |
| Anti-mouse CD150 Monoclonal Antibody PE-Dazzle 594 (TC15-12F12.2) | BioLegend | Cat# 115936 RRID: AB_2565961 |
| Anti-mouse CD16/CD32 Monoclonal Antibody Brilliant Violet 711 (93) | BioLegend | Cat# 101337 RRID: AB_2565637 |

| | | |
|---|---|---|
| Anti-mouse Flt-3 Monoclonal Antibody APC (A2F10) | BioLegend | Cat# 135310<br>RRID: AB_2107050 |
| Anti-mouse CD34 Monoclonal Antibody PE (MEC 14.7) | Thermo Fisher Scientific | Cat# RM3604<br>RRID: AB_10376011 |
| Anti-puromycin Alexa Fluor 647 | Arguello lab | |
| Anti-puromycin Alexa Fluor 488 | Arguello lab | |
| Normal Rat Serum | Thermo Fisher | Cat# No.10710C |
| Bacterial and virus strains | | |
| Biological samples | | |
| Chemicals, peptides, and recombinant proteins | | |
| Cisplatin | Sigma-Aldrich | Cat# P439425MG |
| 191/193Ir DNA Intercalator | Fluidigm | Cat# 201192B |
| Paraformaldehyde | VWR | Cat# PI28908 |
| Ethylenediaminetetraacetic acid (EDTA) | Fisher Scientific | Cat# BP118500 |
| Maxpar 10x Barcode Perm Buffer | Fluidigm | Cat# 201057 |
| 2-Deoxy-D-Glucose | Sigma-Aldrich | Cat# D6134 |
| Oligomycin | Sigma-Aldrich | Cat# 75351 |
| Puromycin | Sigma-Aldrich | Cat# P7255 |
| Critical commercial assays | | |
| LIVE/DEAD™ Fixable Violet Dead Cell Stain Kit, for 405 nm excitation | Thermo Fisher | Cat# L34963 |
| FoxP3 Transcription Factor Staining Buffer Kit | Tonbo Biosciences | Cat# TNB-0607-KIT |
| EQ$^{TM}$ Four Element Calibration Beads | Fluidigm | Cat# 201078 |
| ZOMBIE NIR | Thermo Fisher | WAITING |
| Deposited data | | |

| | | |
|---|---|---|
| Experimental models: cell lines | | |
| MOLT3 (human, male) | Lab of Jeroen Roose | N/A |
| JURKAT (human, male) | Lab of Jeroen Roose | N/A |
| HUT78 (human, male) | Lab of Jeroen Roose | N/A |
| CCRF-CEM (human, female) | Lab of Jeroen Roose | N/A |
| Experimental models: organisms/strains | | |
| Mouse C57BL/6J | Jeroen P. Roose Laboratory | N/A |
| Mouse RoLoRiG | Jeroen P. Roose Laboratory | Oncogene 2020 |
| B6.129S4-Krastm4Tyj/J (KRas$^{G12D}$ mice) | The Jackson Laboratory | Stock no.008179 |
| Mouse Mx1CRE | Passague Laboratory | N/A |
| Oligonucleotides | | |
| Primers | | |
| Recombinant DNA | | |
| Software and algorithms | | |
| PhenoGraph | (Levine et al., 2015) | https://doi.org/10.1016/j.cell.2015.05.047 |
| Umap | | |
| GraphPad prism 9 | GraphPad Software | http://www.graphpad.com |
| Cytobank | | |
| Python | | |
| Other | | |
| Affinity Designer | | |
| FlowJo v10 | | |

*LIFE SCIENCE TABLE WITH EXAMPLES FOR AUTHOR REFERENCE*

| REAGENT or RESOURCE | SOURCE | IDENTIFIER |
|---|---|---|
| Antibodies | | |
| Bacterial and virus strains | | |
| Biological samples | | |
| Chemicals, peptides, and recombinant proteins | | |
| Critical commercial assays | | |
| EasyTag EXPRESS 35S Protein Labeling Kit | PerkinElmer | NEG772014MC |
| CaspaseGlo 3/7 | Promega | G8090 |
| TruSeq ChIP Sample Prep Kit | Illumina | IP-202-1012 |
| Deposited data | | |
| Raw and analyzed data | This paper | GEO: GSE63473 |
| B-RAF RBD (apo) structure | This paper | PDB: 5J17 |
| Human reference genome NCBI build 37, GRCh37 | Genome Reference Consortium | http://www.ncbi.nlm.nih.gov/projects/genome/assembly/grc/human/ |
| Nanog STILT inference | This paper; Mendeley Data | http://dx.doi.org/10.17632/wx6s4mj7s8.2 |
| Affinity-based mass spectrometry performed with 57 genes | This paper; Mendeley Data | Table S8; http://dx.doi.org/10.17632/5hvpvspw82.1 |
| Experimental models: cell lines | | |
| Experimental models: organisms/strains | | |
| *D. melanogaster*: RNAi of Sxl: y[1] sc[*] v[1]; P{TRiP.HMS00609}attP2 | | |
| Mouse: R6/2: B6CBA-Tg(HDexon1)62Gpb/3J | The Jackson Laboratory | JAX: 006494 |
| Mouse: | | |
| Oligonucleotides | | |
| | This paper | N/A |

| | | |
|---|---|---|
| Primers | This paper | N/A |
| | This paper | N/A |
| Recombinant DNA | | |
| | Clonetech | Cat#632162 |
| | This paper | N/A |
| | Drosophila Genomics Resource Center | DGRC:5666; FlyBase:FBcl0130415 |
| | Chen et al., 2013 | N/A |
| | Thoreen et al., 2009 | Addgene Plasmid #21339 |
| Software and algorithms | | |
| Other | | |

**_PHYSICAL SCIENCE TABLE WITH EXAMPLES FOR AUTHOR REFERENCE_**

| REAGENT or RESOURCE | SOURCE | IDENTIFIER |
|---|---|---|
| Chemicals, peptides, and recombinant proteins | | |
| QD605 streptavidin conjugated quantum dot | Thermo Fisher Scientific | Cat#Q10101MP |
| Platinum black | Sigma-Aldrich | Cat#205915 |
| Sodium formate BioUltra, ≥99.0% (NT) | Sigma-Aldrich | Cat#71359 |
| Chloramphenicol | Sigma-Aldrich | Cat#C0378 |
| Carbon dioxide ($^{13}C$, 99%) (<2% $^{18}O$) | Cambridge Isotope Laboratories | CLM-185-5 |
| Poly(vinylidene fluoride-co-hexafluoropropylene) | Sigma-Aldrich | 427179 |
| PTFE Hydrophilic Membrane Filters, 0.22 µm, 90 mm | Scientificfilters.com/Tisch Scientific | SF13842 |
| Critical commercial assays | | |
| Folic Acid (FA) ELISA kit | Alpha Diagnostic International | Cat# 0365-0B9 |
| TMT10plex Isobaric Label Reagent Set | Thermo Fisher | A37725 |
| Surface Plasmon Resonance CM5 kit | GE Healthcare | Cat#29104988 |
| NanoBRET Target Engagement K-5 kit | Promega | Cat#N2500 |
| Deposited data | | |
| B-RAF RBD (apo) structure | This paper | PDB: 5J17 |
| Structure of compound 5 | This paper; Cambridge Crystallographic Data Center | CCDC: 2016466 |
| Code for constraints-based modeling and analysis of autotrophic *E. coli* | This paper | https://gitlab.com/elad.noor/sloppy/tree/master/rubisco |
| Software and algorithms | | |
| Gaussian09 | Frish et al., 2013 | https://gaussian.com |
| Python version 2.7 | Python Software Foundation | https://www.python.org |
| ChemDraw Professional 18.0 | PerkinElmer | https://www.perkinelmer.com/category/chemdraw |
| Weighted Maximal Information Component Analysis v0.9 | Rau et al., 2013 | https://github.com/ChristophRau/wMICA |
| Other | | |
| DASGIP MX4/4 Gas Mixing Module for 4 Vessels with a Mass Flow Controller | Eppendorf | Cat#76DGMX44 |
| Agilent 1200 series HPLC | Agilent Technologies | https://www.agilent.com/en/products/liquid-chromatography |
| PHI Quantera II XPS | ULVAC-PHI, Inc. | https://www.ulvac-phi.com/en/products/xps/phi-quantera-ii/ |
